# Increased Neurite Orientation-Dispersion and Density in the TgCRND8 Mouse Model of Amyloidosis: Inverse Relation with Functional Connectome Clustering and Modulation by Interleukin-6

**DOI:** 10.1101/562348

**Authors:** Luis M. Colon-Perez, Kristen R. Ibanez, Mallory Suarez, Kristin Torroella, Kelly Acuna, Edward Ofori, Yona Levites, David E. Vaillancourt, Todd E. Golde, Paramita Chakrabarty, Marcelo Febo

## Abstract

Extracellular β-amyloid (Aβ) plaque deposits and inflammatory immune activation are thought to alter various aspects of tissue microstructure, such as extracellular free water, fractional anisotropy and diffusivity, as well as the density and geometric arrangement of axonal processes. Quantifying these microstructural changes in Alzheimer’s disease and related neurodegenerative dementias could serve to accurately monitor or predict disease course. In the present study we used high-field diffusion magnetic resonance imaging (dMRI) to determine how Aβ and inflammatory interleukin-6 (IL6), alone or in combination, affect *in vivo* tissue microstructure in the TgCRND8 mouse model of Alzheimer’s-type Aβ deposition. TgCRND8 and non-transgenic (nTg) mice expressing brain-targeted IL6 or enhanced glial fibrillary protein (EGFP controls) were scanned at 8 months of age using a 2-shell, 54-gradient direction dMRI sequence at 11.1 Tesla. Images were processed using the free water elimination method and the neurite orientation dispersion and density imaging (NODDI) model. DTI and NODDI processing in TgCRND8 mice revealed a microstructure pattern consistent with reduced white matter integrity along with an increase in density and geometric complexity of axonal and dendritic processes. This included reduced FA, mean diffusivity (MD), and free water (FW), and increased ‘neurite’ density (NDI) and orientation dispersion (ODI). IL6 produced a ‘protective-like’ effect on FA in TgCRND8 mice, although there were minimal microstructure changes in these mice compared IL6 expressing nTg mice. In addition, we found that NDI and ODI had an inverse relationship with the functional connectome clustering coefficient, which was affected by Aβ and IL6. The relationship between NODDI and graph theory metrics suggests that increasing the density and orientation dispersion of neurites may relate to diminished functional network organization in the brain.

## INTRODUCTION

Excess production and oligomerization of the β-amyloid (Aβ) protein, the main constituent of amyloid plaques, substantially modifies neuronal morphology (Biscaro et al., 2009; Spires and Hyman, 2004). Important, Aβ alters neuronal homeostasis and synaptic plasticity by modifying the extracellular environment of neuronal soma, axons, and synapses (Mueggler et al., 2004; Sykova et al., 2005). Aβ is implicated in the formation of dystrophic ‘neurites’ (Sadleir et al., 2016; Zhang et al., 2009), deficits in synaptic plasticity and excitability (Cummings et al., 2015; Varga et al., 2015), cell membrane protein aggregation (Shrivastava et al., 2017), reduced integrity of major fiber tracts (Mayo et al., 2017; O’Dwyer et al., 2011; Teipel et al., 2014) and impaired activity in resting state networks (Hedden et al., 2009; Jones et al., 2016). In addition, Aβ is immunogenic (Dalgediene et al., 2013), leading to the increased activated microglia in the brain (Mirzaei et al., 2016; Serriere et al., 2015). Positron emission tomography (PET) studies using microglia- or Aβ-specific radiotracers provide compelling evidence of an overlapping presence of activated microglia and amyloid burden in Alzheimer’s disease (Kreisl et al., 2013; Liu et al., 2015; Mirzaei et al., 2016; Parbo et al., 2017; Schuitemaker et al., 2013). For instance, a longitudinal study identified a transient elevation in microglia early in mild cognitive impairment (MCI) which peaked later in Alzheimer’s disease (Fan et al., 2017). Aged transgenic mice expressing familial Alzheimer’s mutations to amyloid precursor protein and presenilin-1 with deletion of exon-9 (APP/PS1dE9) had higher *in vivo* cortical and hippocampal accumulation of an activated microglial-targeted radiotracer, 18kD translocator protein (TSPO), than aged wildtype or young Tg mice (Liu et al., 2015). Taken together, the above findings strongly suggest that accrual of Aβ plaques and concomitant reactive microgliosis may underlie pathologic dysfunction in Alzheimer’s by causing deleterious changes to brain parenchymal microstructure. In this study, we asked the question whether we can utilize high field diffusion magnetic resonance imaging (dMRI) to quantitatively assess amyloid pathology and immune-related mechanisms in a mouse model of Alzheimer’s disease and establish their mechanistic links with intrinsic brain activity measured by resting state functional MRI (fMRI)(Brown et al., 2018).

Diffusion tensor imaging (DTI) offers a powerful approach to generate metrics that describe the movement of water in brain tissue compartments. Advanced methods for the modeling of tissue diffusivity beyond standard DTI processing schemes may provide additional insight on the effects of pathogenic Aβ and immune mechanisms in Alzheimer’s disease. For example, increased radial diffusivity was associated with worsening of dementia symptoms in dMRI scans processed using a two-compartment model of diffusion known as the free water imaging method (Ji et al., 2017; Maier-Hein et al., 2015). Also, a multi-compartment diffusion model termed ‘Neurite Orientation Dispersion and Density Imaging’ (NODDI) revealed significant changes in NODDI metrics in young onset Alzheimer’s disease (Parker et al., 2018; Slattery et al., 2017), and correlations with cortical tau immunoreactivity in the rTg4510 mouse tauopathy model of frontal temporal dementia (Colgan et al., 2016). The free water method (Pasternak et al., 2009) provides quantitative maps that have been associated with neuroinflammatory and neurodegenerative mechanisms (Ofori et al., 2015; Pasternak et al., 2012; Pasternak et al., 2015), and the NODDI model provides quantitative maps associated with axonal and dendritic dispersion and density (Colgan et al., 2016; Grussu et al., 2017; Zhang et al., 2012). Despite the advances in dMRI processing, preclinical high field imaging studies in mouse models are needed to determine the changes in brain microstructure that link to Aβ or neuroinflammation, and the impact of such changes on functional network activity.

In the present study we investigated the effects of Aβ on diffusion MRI-derived NODDI and free water metrics in transgenic (Tg)CRND8 mice with early onset amyloidosis (Chishti et al., 2001). To mimic the effects of chronic microgliosis and astrogliosis, we additionally used adeno-associated viral vector (AAV)-mediated brain-targeted expression of an inflammatory cytokine, Interleukin (IL)-6. IL6 expression results in widespread gliosis in the CNS in the absence of overt amyloidogenic or neurodegenerative pathology (Chakrabarty et al., 2010b), and was thus considered ideal for testing the individual or combined effects of Aβ and neuroinflammation on *in vivo* tissue microstructure in mouse brain. Collectively, free water and NODDI processing in Aβ-expressing TgCRND8 mice revealed a microstructure pattern consistent with a reduced volume of the extracellular space along with an increase in geometrically misoriented (dispersed) axons or dendritic processes. Interestingly, we discovered a novel relationship between NODDI metrics modeling the density and orientation dispersion of neurites and a measure of functional network clustering that was similarly affected by Aβ and IL6.

## MATERIALS AND METHODS

### Subjects

Mice were housed in sex- and aged-matched groups of 3-4 in conventional cages (29 x 18 x 13 cm) at 20-26°C (lights on from 0700-1900 hours) with food and water *ad libitum*. TgCRND8 mice were maintained in-house by breeding amyloid precursor protein (APP) transgenic (Tg) males (carrying the wild-type retinal degeneration gene) with C57B6/C3H F1 females (Envigo) (Janus et al., 2000). TgCRND8 mice have early onset expression of human mutant APP (Swedish APP KM670/671NL and Indiana APP V717F), which increases human APP 5-times above endogenous murine APP. These mice have aggressive cognitive impairment, Aβ plaque deposits and increased inflammation at 3-4 months of age, synaptic deficits and some synaptic and neuronal loss in hippocampus by 6 mo. (Chishti et al., 2001). In this study, we used 8 month old mice (non-transgenic (nTg) and TgCRND8), an age where there is high levels of forebrain amyloidosis. All groups were sex and age-matched. All procedures received prior approval from the Institutional Animal Care and Use Committee of the University of Florida and follow all applicable NIH guidelines.

### Adeno-associated viral vector (AAV) preparation and brain delivery

Recombinant adeno-associated virus subtype 2/1 (rAAV2/1) for intracerebroventricular (ICV) injections expressing murine IL6 (m-IL6) or enhanced green fluorescent protein (EGFP) under the control of the cytomegalovirus enhancer/chicken β actin promoter were generated as described previously (Chakrabarty et al., 2010b). Briefly, rAAV vectors expressing IL6 under the control of the cytomegalovirus enhancer/chicken beta actin (CBA) promoter, a WPRE, and the bovine growth hormone polyA were generated by plasmid transfection with helper plasmids in HEK293T cells. Forty-eight hours after transfection, cells were harvested and lysed in the presence of 0.5% Sodium Deoxycholate and 50U/ml Benzonase (Sigma) by freeze thawing and the virus isolated using a discontinuous Iodixanol gradient and affinity purified on a HiTrap HQ column (Amersham). The genomic titer of each virus was determined by quantitative polymerase chain reaction (qPCR). TgCRND8 and nTg mice were injected with 2 μl of rAAV (~2E10 viral genomes) into cerebral ventricles using a 10 μl Hamilton syringe with a 30 g needle on day P0 as described before (Chakrabarty et al., 2010b; Chakrabarty et al., 2015). Animals were euthanized within 1-2 days of imaging.

### Experimental groups

The following experimental groups were included in this study: nTg-Controls expressing AAV-EGFP (n = 14), nTg-IL6 expressing IL6 (n = 14), TgCRND8-Controls expressing AAV-EGFP (n = 10) and TgCRND8-IL6 expressing IL6 (n = 10). All 4 groups were sex-balanced. Statistical analysis showed no sex differences, thus data for male and female mice were grouped. One dMRI scan in the nTg-IL6 group had excess, uncorrectable noise and was removed (leaving 13 subjects in this group and a total of 47 mice for the entire study).

### MRI acquisition

Mouse brain images were acquired on an 11.1 Tesla MRI scanner (Magnex Scientific Ltd., Oxford, UK) with a Resonance Research Inc. gradient set (RRI BFG-113/60, maximum gradient strength of 1500 mT/m at 150 Amps and a 130 μs risetime; RRI, Billerica, MA) and controlled by a Bruker Paravision 6.01 console (Bruker BioSpin, Billerica, MA). An in-house 2.0 x 3.5 cm quadrature surface transmit/receive coil tuned to 470.7MHz (^1^H resonance) was used for B1 excitation and signal detection (AMRIS Facility, Gainesville, FL). Anesthesia was induced under 1.5-2.0% isoflurane (0.1-0.15 L/min) delivered in medical grade air (70%N_2_/30% O_2_) and levels were then kept at 1% through functional neuroimaging and then increased to 1.5% during diffusion MRI to maintain a slower breathing rate during the latter acquisition. Mice were placed on a custom plastic bed with a respiratory pad underneath the abdomen. Respiratory rates were monitored continuously, and body temperature was maintained at 37-38°C using a warm water recirculation system (SA Instruments, Inc., New York).

A T2-weighted Turbo Rapid Acquisition with Refocused Echoes (RARE) sequence was used for acquiring anatomical scans with the following parameters: echo time (TE) = 41 ms, repetition time (TR) = 4 seconds, RARE factor = 16, number of averages = 12, field of view (FOV) 15 mm x 11 mm and 0.7 mm thick slice, and 256 x 256 x 17 coronal (axial) slices covering the entire brain from the rostral-most extent of the prefrontal cortical region and caudally to the upper brainstem/cerebellum.

Functional images were collected using a single-shot spin-echo echo planar imaging (EPI) sequence with the following parameters: TE = 15 ms, TR = 2 seconds, 180 repetitions, 15 x 11 mm in plane, 14 slices with 0.9 mm thickness per slice, data matrix = 64 x 48. No stimuli were presented during functional scanning and respiratory rates, isoflurane concentration, body temperature, lighting and room conditions were kept constant across subjects.

Diffusion weighted scans were acquired using a 4-shot, 2-shell spin echo planar diffusion imaging (DTI EPI) sequence in Bruker Paravision, with TR = 4 seconds, TE = 19 ms, number of averages = 4, gradient duration δ = 3 ms, diffusion time Δ = 8 ms, 54 images with 3 different diffusion weightings, two b=0, 6 directions with b=600 s/mm^2^, and 46 directions with b=2000 s/mm^2^. Image saturation bands were placed on either side and below the brain to suppress non-brain signal during image acquisition. Diffusion images had the same FOV and slice thickness as the anatomical scan but with a lower resolution data matrix size of 128 x 96 and 17 slices (resolution: 0.117 mm x 0.117 mm x 0.7 mm) in the same space as anatomical scans. This allowed careful manual outlining of regions to be analyzed by using T2 scans and diffusion maps (see below).

### Brain region of interest (ROI) segmentation for dMRI scans

ITKSNAP (Yushkevich et al., 2006) was used to manually segment hippocampus and WM areas such as anterior commissure, fimbria, genu and splenium of the corpus callosum, and cerebral peduncle (Figure 1, top panel). We selected these axonal pathways because of their critical roles in carrying information between nuclei in striatum and forebrain, prefrontal cortical areas, sensorimotor, temporal, parietal and retrosplenial cortical structures, and the hippocampal-septal system. The T2 high resolution anatomical and the fractional anisotropy (FA) maps were used as guides to determine boundaries between tissue structures. Every effort was made to avoid crossing boundaries between grey and white matter and between the latter two tissue types with the cerebrospinal fluid (CSF) filled lateral ventricles. We sampled the lateral cerebral ventricles to corroborate upper values of diffusivity in a region with unrestricted water mobility.

**Figure 1.**
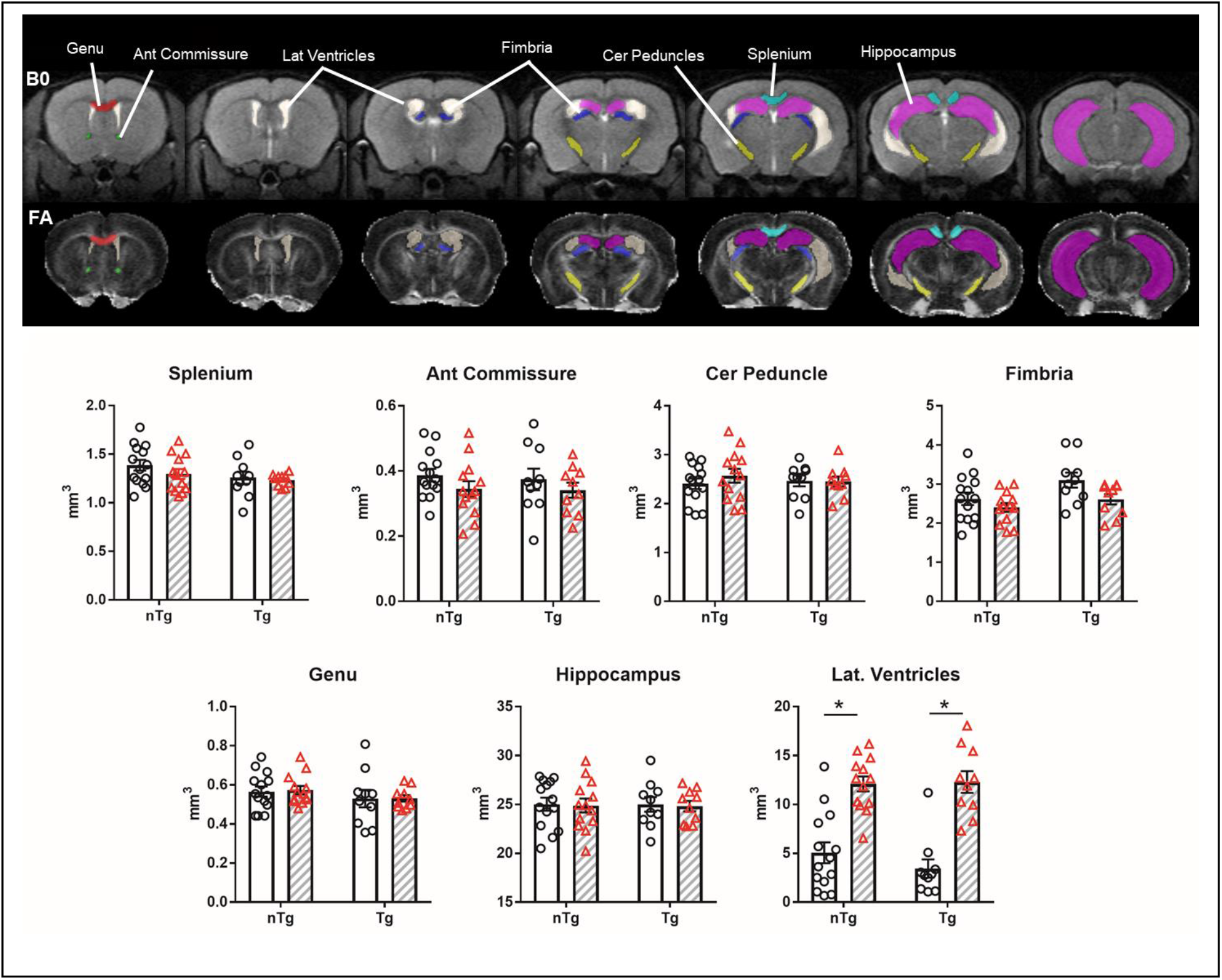
Proinflammatory IL6 increases lateral ventricle volume in Tg and nTg mice. No additional volumetric differences were observed in WM and hippocampus. Top, Segmented ROIs are overlaid on representative B0 and FA maps and ROI names are indicated on the B0 map. Bottom, WM and hippocampus volumes (in mm^3^). Clear bars (and circles) are control groups and hashed bars (and triangles) are IL6 treated mice. *significant difference between control and IL6 (Tukey’s post hoc test, p<0.05). All data are mean ± standard error.

### Diffusion MRI processing

Diffusion MRI scans were processed using tools available on FMRIB software library – FSL (Smith et al., 2004). Diffusion images were first screened for low signal-to-noise (SNR) slices in the image series. Images with excess motion, or excessively low SNR (images with mean % attenuation level below that for a given b-value, EPI ghosting, or images with excess gradient-associated noise), were removed from the image series and gradient b-vector files corrected by removing gradient directions for the removed images. Eddy correction was used for adjusting slight movements during image acquisition and gradient files rotated according to the motion correction vectors.

### Free water imaging

The free water elimination procedure was carried out prior to tensor element construction and calculation of FA and mean diffusivity (MD) maps using the DTIFIT program in FSL (Jenkinson et al., 2012). The free water elimination method uses the Beltrami regularization technique to fit attenuation-normalized diffusion datasets to a bi-exponential model of water diffusion, which estimates the fractional contribution of freely diffusing unrestricted water to the overall signal per voxel (Pasternak et al., 2009). The free water elimination algorithm is implemented in MATLAB (Natick, MA) using custom code (provided by Dr. Ofer Pasternak, Harvard Medical School) and modified for use with mouse brain diffusion MRI (DeSimone et al., 2017). The code generates a mouse brain image of free water (indexed from 0 to 1, the latter meaning highest extracellular free water) and a diffusion tensor with free water fraction removed (free water corrected tensor). The corrected tensor was processed using math tools on FSL to produce maps of FA, MD, axial and radial diffusivities (AD and RD, respectively).

### NODDI processing

We analyzed dMRI scans following processing with the NODDI model (Zhang et al., 2012). This was implemented within the <u>A</u>ccelerated <u>M</u>icrostructure <u>I</u>maging via <u>C</u>onvex <u>O</u>ptimization (AMICO) framework (Daducci et al., 2015), which accelerates the model fitting procedure in standard Linux PC’s running Python 2.7. NODDI incorporates a multi-compartment model of the diffusion signal. The model includes an isotropic fraction of free unrestricted and hindered water (e.g., cerebral spinal fluid) and an anisotropic model that includes an intracellular fraction with zero radius infinite cylinders (high diffusion coefficients along the length of the principal axis only) and an extracellular fraction with non-zero diffusion perpendicular to the principal axis. The latter adds morphological information to the tissue model of the extracellular environment surrounding axons and dendrites (occupied by neurons and non-neuronal cell bodies) (Zhang et al., 2012). The fitting of the intracellular volume fraction (i.e., zero-radius cylinders or ‘sticks’) involves the use of a Bingham-Watson series used in statistical shape analysis (Maier-Hein et al., 2015) and the results of the fitting process provides an index of the degree of orientation dispersion of so-called ‘neurites’. For the parameter fitting procedure, we assumed a value of *in vivo* intrinsic diffusivity of 1.7 μm^2^/ms and isotropic diffusivity of 3.0 μm^2^/ms (Grussu et al., 2017). The whole brain model-fitting generates maps of neurite density (NDI: relative concentration of zero-radius cylinders), orientation dispersion (ODI: 0 for no dispersion as in highly organized parallel fiber bundles to a maximum of 1 for the highest degree of dispersion, as in cerebral cortical grey matter), and the isotropic free water fraction (ISO: 0 low isotropy to a maximum isotropy of 1) (Grussu et al., 2017).

### Resting state fMRI processing

Binary masks outlining mouse brain boundaries were automatically generated on MATLAB using Pulsed Coupled Neural Networks (PCNN3D) on high resolution T2 Turbo-RARE anatomical scans (Chou et al., 2011). The binary ‘brain-only’ masks were multiplied by T2 anatomical scans to null voxels outside the brain. This step was found to enhance the precision of subject-to-atlas linear registration. The cropped brain images are aligned to a mouse brain template using the FSL program FLIRT (Jenkinson et al., 2002) (Supplemental Figure 1). Registration matrices for each subject are saved and used to transform functional datasets to template space for preprocessing and analysis (Supplemental Figure 1). Resting state processing steps include: (1) removal of time series spikes, (2) slice timing correction, (3) motion correction, (4) subject-to-atlas registration, (5) regression of movement, white matter and CSF (ventricle) signals, (6) bandpass filtering (0.01-0.1Hz), and (7) spatial blurring (0.6 mm FWHM). Average BOLD signals from each region of interest (ROI) were extracted based on their atlas location and used in voxel-wise cross correlation to generate Pearson correlation maps and 4005 pairwise correlation coefficients to be imported to MATLAB as edge weights for network analysis. All correlation r values are z transformed prior to statistical analyses. For seed-based resting state fMRI analysis of cortical and memory structures, connectivity maps of somatosensory cortex, hippocampus, and entorhinal cortex were generated for each subject (Supplemental Figure 2) and analyzed using a multivariate modeling approach in Analysis of Functional Neuroimages (AFNI 3dMVM)(Cox, 1996). All group statistical maps for main effects and full factorial interactions were FDR corrected (Chen et al., 2014).

### Network analyses

Functional network graphs are analyzed using Brain Connectivity Toolbox (Rubinov and Sporns, 2010). Graphs with 4,005 entries are organized in MATLAB [graph size = n(n-1)/2, where n is the number of nodes represented in the graph, or 90 ROI]. The z-scores are thresholded to equalize density prior to analyses (multiple node density thresholds from 2-20% are applied). Edge weights are normalized between 0 to 1. Node strength (sum of edge weights), clustering coefficient (the degree to which nodes cluster together in groups), average path length (the potential for communication between pairs of structures), modularity (quantifies the degree to which the network may be subdivided into such clearly delineated groups or communities), and small worldness (the degree to which mouse functional brain networks under study deviate from randomly connected networks) are calculated for unweighted graphs and statistically compared between groups (Boccaletti et al., 2006; Humphries and Gurney, 2008; Newman and Girvan, 2004; Newman, 2003; Saramaki et al., 2007). Brain networks are visualized using BrainNet Viewer (Xia et al., 2013). The 3D networks are generated with undirected edges weights ≥ 0.2. In these brain networks, the node size and color are scaled by the node strength, and edges are scaled by z-scores.

### Preparation of brain tissue for immunohistochemistry and ELISA

Following euthanasia, the right brains was fixed in 10% normal buffered formalin and the left brains were snap frozen. Formalin fixed brains were paraffin embedded and analyzed using the following antibodies according to immunohistochemistry protocols described previously (Chakrabarty, 2010; Chakrabarty, 2015): Glial Fibrillary Acidic Protein (GFAP, Cell Signaling, 1:1000) and ionized calcium binding adaptor molecule 1 (Iba1, Wako, 1:1000). Immunohistochemically stained sections were captured using the Scanscope XT image scanner (Aperio) and analyzed using the ImageScope program. Brightness and contrast alterations were applied identically on captured images using Adobe Photoshop CS3. Snap-frozen forebrains (without olfactory bulb and cerebellum) were homogenized in RIPA Buffer (Boston Bioproducts) and the homogenate was centrifuged at 100,000 x g for 1 h at 4°C. Protein concentration in the supernatant was determined using BCA Protein Assay kit (Pierce). mIL-6 levels were evaluated using sandwich capture ELISA assays using the RIPA soluble mouse forebrain lysates as per manufacturer’s instructions (BD OptiEIA, BD Biosciences).

### Statistical analyses

For dMRI, manual segmentation masks were used to import to MATLAB the average values for FA, MD, AD, RD (all diffusivities in units of mm^2^/sec), ODI, NDI, and ISO for 7 ROI’s. Diffusion metrics were analyzed in MATLAB (Natick, MA) using a two-factor analysis of variance (ANOVA: Strain [2 levels: nTg and Tg] x Treatment [2 levels: control and IL6]). Tukey’s multiple comparison test was used to assess specific differences between group means. Unless otherwise noted, Tukey’s post hoc test results are reported in figure legends, whereas full factorial ANOVA results are provided in the results sections.

For fMRI, mouse brain atlas-based segmentation masks were used to import to MATLAB the average values for small world coefficient, global node strength, clustering coefficient, modularity, and mean path length. For these measures, we used a two-factor ANOVA (Strain x Treatment) and Tukey’s posthoc test. Statistical analyses of correlation coefficient values per ROI pairs were not carried out on exported values, as these were analyzed on somatosensory, hippocampal and entorhinal cortex maps using ANOVA computations on AFNI 3dMVM (see above). Main effects of strain and treatments and their interactions were statistically assessed.

As an additional step to supplement seed-based and network analyses, we conducted analyses using probabilistic independent component analysis (ICA) in FSL MELODIC (multivariate exploratory linear decomposition into independent components) for 48 resting state fMRI mouse brain scans (Beckmann and Smith, 2004). Prior to ICA, post-processed and atlas-aligned functional scans (see above) were cropped using manually segmented masks. This provided optimal co-registration of fMRI scans with each subject anatomical and with the mouse brain template. In MELODIC, the following steps were carried out: voxel-wise de-meaning, variance normalization, pre-whitening, and data were then projected into a 30-dimensional subspace using PCA, and then ICA estimation was conducted in concatenated maps. Mixture model fitting, rescaling and thresholding was carried out on z-transformed ICA maps (alternative hypothesis test set at p > 0.5). ICA maps likely representing noise or not deemed to include well-defined regions were removed and the rest were considered of interest. The latter were assembled for visualization using slices summary and overlay scripts in FSL, using a lower z threshold of 2.3. Using the Glm tool in FSL a general linear model design matrix was generated to test for main effects of strain, treatment, and strain x treatment interactions, and used in dual regression analysis and randomize permutation testing in FSL to test for differences between groups (Nickerson et al., 2017).

## RESULTS

### IL6 treatment increased lateral ventricle volume

To understand how inflammatory conditioning alters brain connectivity in an Alzheimer’s disease mouse model, we delivered AAV-IL6 into the cerebral ventricles of neonatal TgCRND8. We have previously demonstrated the utility of similar paradigms to examine pathological alterations in Alzheimer’s disease mouse models (Chakrabarty, 2010; Chakrabarty, 2015). Mice were aged to 8 months, imaged and euthanized to confirm the histopathological effects of intracranial expression of IL6 in this study. We observed robust upregulation of astrogliosis (GFAP immunohistochemistry) and microgliosis (Iba-1 immunohistochemistry) in all areas of the brain examined, e.g., forebrain (cortex and hippocampus), basal ganglia (striatum) and midbrain (Supplementary Figure 3A-D). IL-6 protein levels were also upregulated in the whole brain lysates (p<0.01; Supplementary Figure 3E).

To corroborate any morphological differences in WM and hippocampus across subjects as a result of IL6 or Aβ, we analyzed ROI volumes (Figure 1). No volumetric differences were observed between nTg and Tg or control and IL6 treatment in any ROI. However, we discovered that elevated brain IL6 significantly increased the volume of the lateral ventricles (main effect IL6 treatment: F_1,43_ = 63.6 p < 0.0001). The enlarged lateral ventricles were observed in nTg and Tg mice given IL6 treatment.

### Opposing effects of Aβ and IL6 on FA in white matter

Figure 2 summarizes the results for FA as a function of strain (nTg x Tg) and IL6 treatment (IL6 x control). Representative mouse brain B0 and FA maps in Figure 2 illustrate WM structures such as the corpus callosum, fimbria, cerebral peduncle, and their directionality, and also the lateral ventricles and the hippocampus. In splenium, anterior commissure, and cerebral peduncle we observed a main effect of Aβ on FA (splenium: main effect strain F_1,43_ = 15.4 p = 0.0003; cerebral peduncle: strain x treatment interaction F_1,43_ = 11.2 p = 0.002; main effect strain F_1,43_ = 16.5 p = 0.0002; anterior commissure: main effect of treatment F_1,43_ = 8.4 p = 0.006). In splenium, fimbria, and cerebral peduncle IL6 increased FA (splenium: main effect treatment F_1,43_ = 8.4 p = 0.006; fimbria: main effect treatment F_1,43_ = 16.3 p = 0.0002; cerebral peduncle: main effect treatment F_1,43_ = 61 p < 0.0001). No significant effects of IL6 or Aβ were observed in genu and hippocampus.

**Figure 2.**
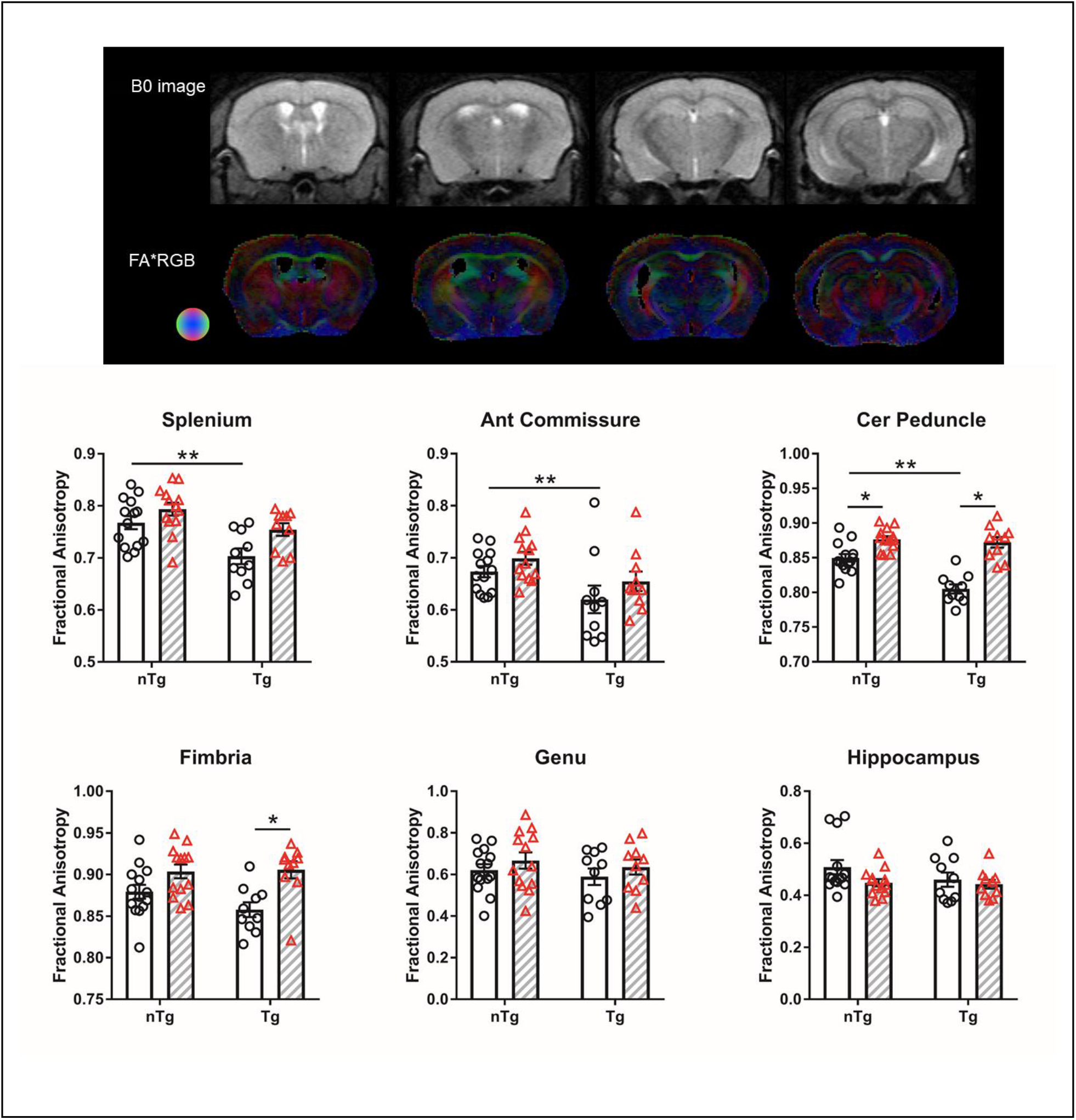
Aβ reduces and IL6 increases FA in WM regions. Representative B0 and FA color map shown. Diffusion direction indicated by colored sphere. Clear bars (and circles) are control groups and hashed bars (and triangles) are IL6 treated. *significant difference between control and IL6 mice; **significant difference between Tg and nTg mice (Tukey’s post hoc test, p<0.05). All data are mean ± standard error.

### Differential effects of Aβ and IL6 on MD, AD and RD metrics in white matter

We observed that increased Aβ in Tg mice was associated with reduced MD in cerebral peduncle (Figure 3; main effect strain F_1,43_ = 6.0 p = 0.01) and genu (main effect strain F_1,43_ = 7.5 p = 0.008). Elevated IL6 expression reduced MD in splenium (main effect treatment F_1,43_ = 10.2 p = 0.002).

**Figure 3.**
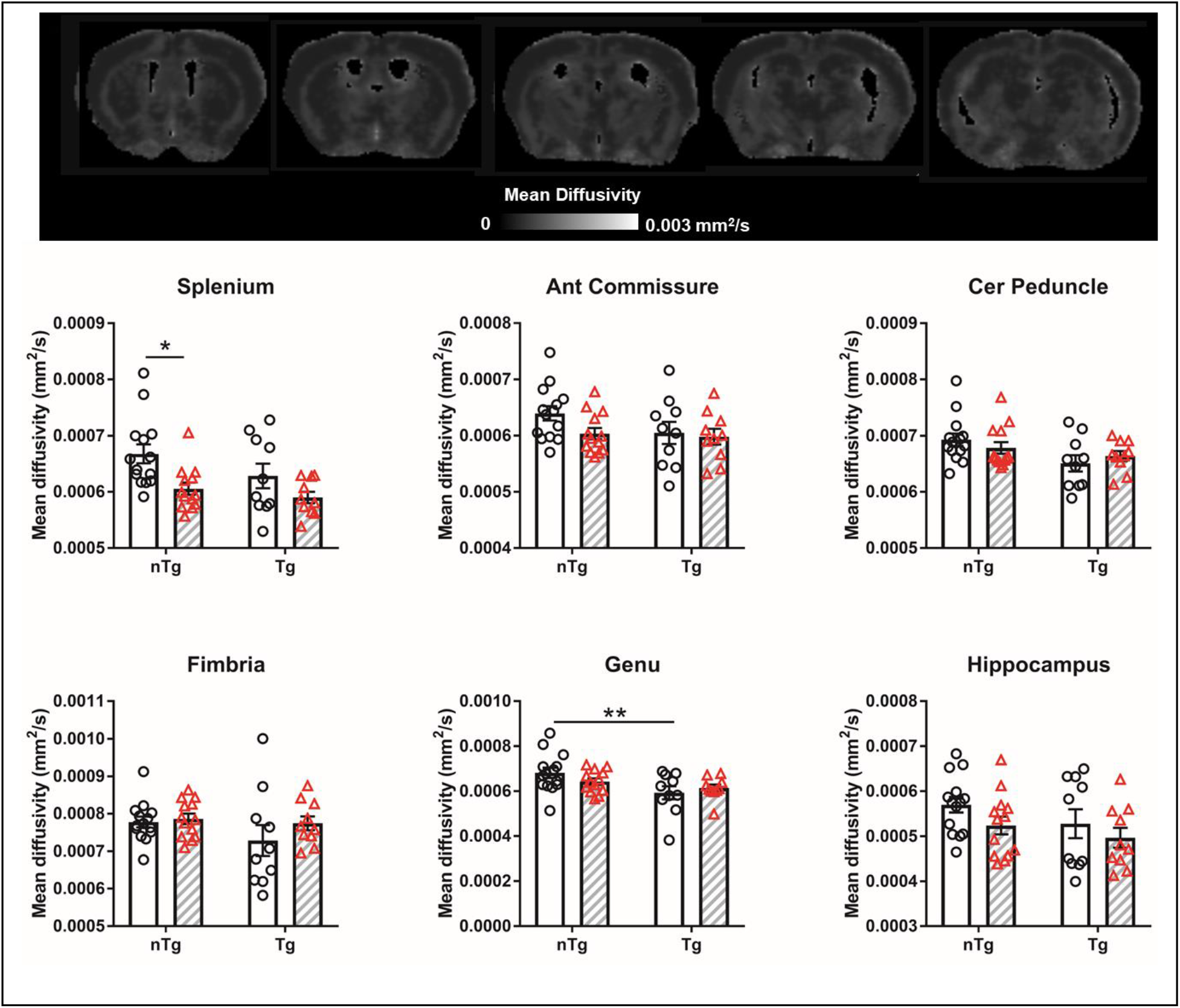
Aβ and IL6 reduce MD in WM regions. Representative MD map shown. Clear bars (and circles) are control groups and hashed bars (and triangles) are IL6 treated. *significant difference between control and IL6 mice; **significant difference between Tg and nTg mice (Tukey’s post hoc test, p<0.05). All data are mean ± standard error.

AD and RD were also assessed (Supplemental Figures 4-5). TgCRND8 mice had lower AD in splenium and cerebral peduncle than nTg (splenium: main effect strain F_1,43_ = 45.2 p < 0.0001; cerebral peduncle: main effect strain F_1,43_ = 18.8 p < 0.0001). In the splenium, IL6 reduced AD in nTg mice (strain x treatment interaction F_1,43_ = 4.4 p = 0.04). In cerebral peduncle and fimbria AD was increased in Tg mice (cerebral peduncle: treatment x strain interaction F_1,43_ = 7.3 p = 0.01 and main effect treatment F_1,43_ = 17.1 p = 0.0002; fimbria: main effect treatment F_1,43_ = 12.5 p = 0.001). TgCRND8 had higher RD in cerebral peduncle than nTg mice (main effect of strain F_1,43_ = 13.6 p = 0.0006). IL6 reduced RD in fimbria and cerebral peduncle of TgCRND8 mice (fimbria: main effect treatment F_1,43_ = 16.8 p = 0.0002; cerebral peduncle: main effect treatment F_1,43_ = 48.5 p < 0.0001) and in the cerebral peduncle of nTg mice (treatment x strain interaction F_1,43_ = 5.3 p = 0.02). No effect of Aβ or IL6 on MD, AD, and RD were observed in genu or hippocampus.

### Aβ and IL6 reduce free water in white matter and hippocampus

We observed that TgCRND8 had lower free water values in fimbria and hippocampus than nTg mice (fimbria: treatment x strain interaction F_1,43_ = 5.0 p = 0.03 and main effect strain F_1,43_ = 7.5 p = 0.009; hippocampus: treatment x strain interaction F_1,43_ = 6.8 p = 0.01 and main effect strain F_1,43_ = 12.9 p = 0.0008) (Figure 4). IL6 reduced free water only in nTg mice in anterior commissure, cerebral peduncle, genu and hippocampus (anterior commissure: main effect strain F_1,43_ = 9.4 p = 0.004; cerebral peduncle: main effect strain F_1,43_ = 18.7 p < 0.0001; genu: main effect strain F_1,43_ = 5.1 p = 0.03; hippocampus: main effect treatment F_1,43_ = 5.8 p = 0.02).

**Figure 4.**
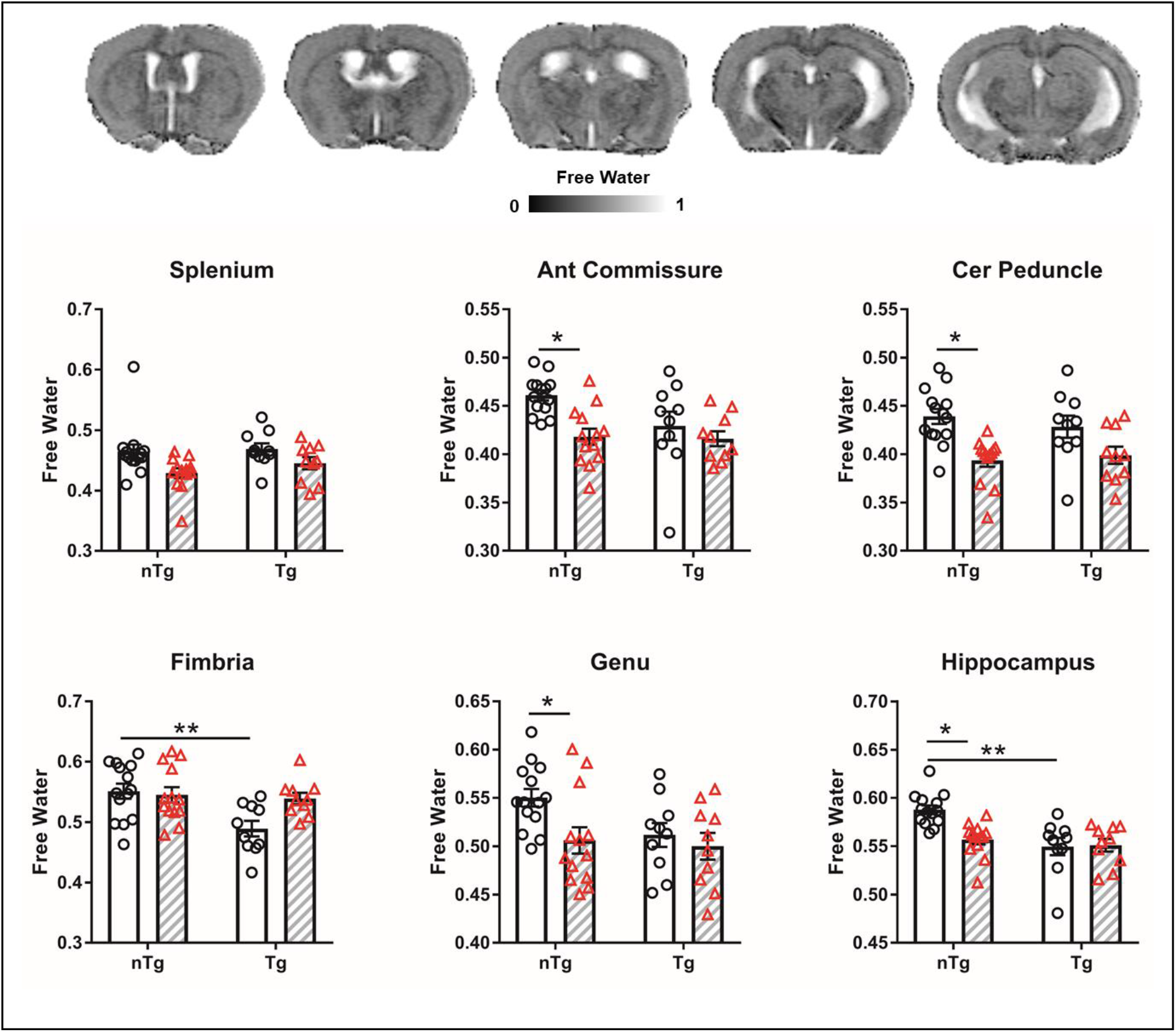
Aβ and IL6 reduce free water in WM regions and hippocampus. Representative free water map shown. Clear bars (and circles) are control groups and hashed bars (and triangles) are IL6 treated. *significant difference between control and IL6 mice; **significant difference between Tg and nTg mice (Tukey’s post hoc test, p<0.05). All data are mean ± standard error.

### Aβ and IL6 increase NDI (NODDI intracellular volume fraction)

We conducted NODDI analysis to examine the whether Aβ and IL6 altered the fraction of tissue corresponding to axons or dendrites (Figure 5). Highest neurite density is observed in large WM tracts such as the corpus callosum (which appear bright), and varying degrees of NDI are observed across grey matter areas (which appear in various light grey tones). TgCRND8 mice had higher NDI values in the cerebral peduncle and hippocampus (cerebral peduncle: main effect strain F_1,43_ = 13.4 p = 0.0007; hippocampus: main effect strain F_1,43_ = 11.8 p = 0.001). IL6 treatment increased NDI values only in nTg mice in the splenium, anterior commissure, cerebral peduncle, genu and hippocampus (splenium: main effect treatment F_1,43_ = 12.9 p = 0.0008; anterior commissure: main effect treatment F_1,43_ = 10.6 p = 0.002; cerebral peduncle: main effect treatment F_1,43_ = 11.1 p = 0.002; genu: main effect treatment F_1,43_ = 9.9 p = 0.003; hippocampus: main effect treatment F_1,43_ = 10.3 p = 0.002).

**Figure 5.**
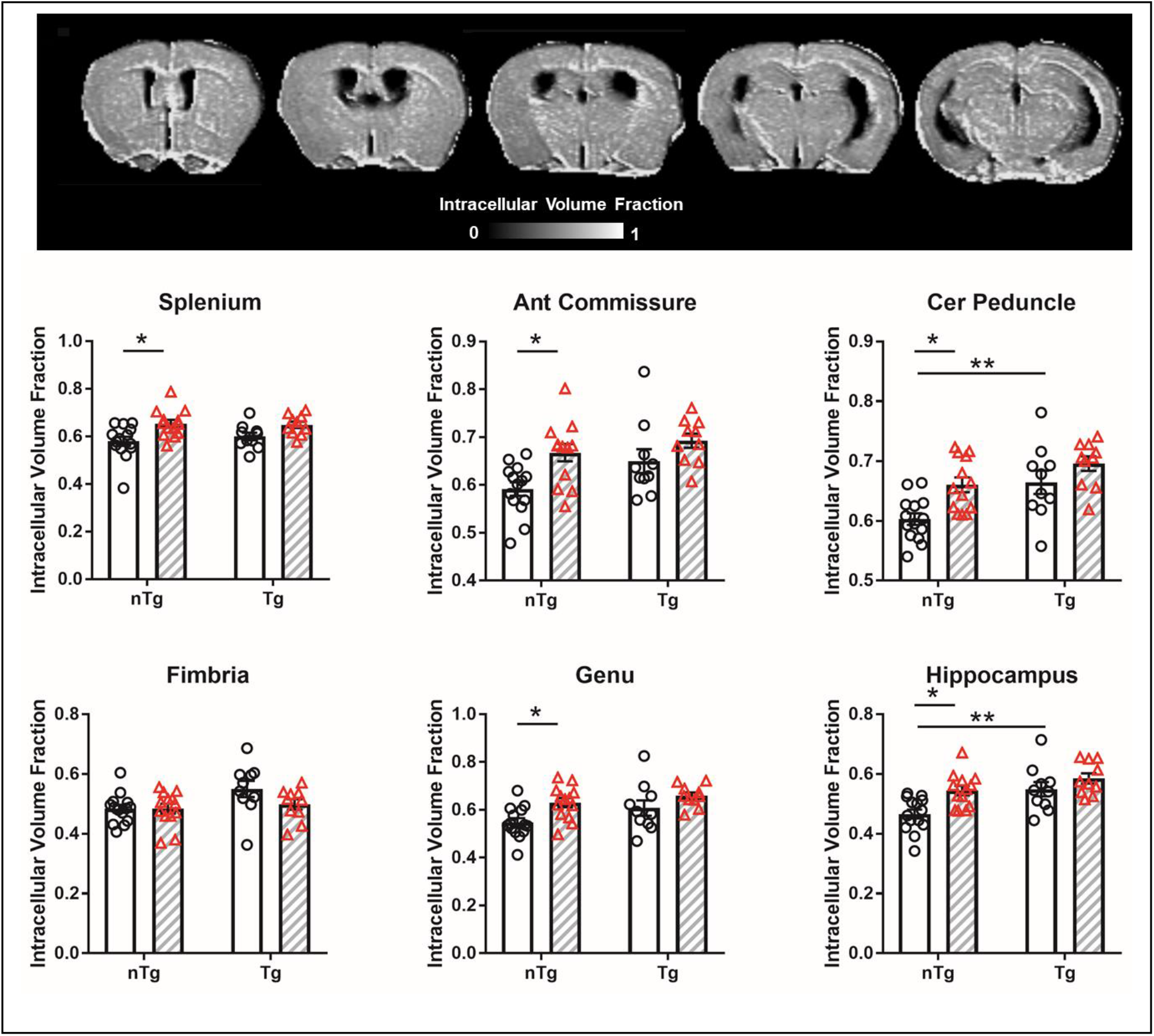
Aβ and IL6 increase NDI in WM regions and hippocampus. The effects of IL6 are observed only in nTg mice. Representative NDI map shown. Clear bars (and circles) are control groups and hashed bars (and triangles) are IL6 treated. *significant difference between control and IL6 mice; **significant difference between Tg and nTg mice (Tukey’s post hoc test, p<0.05). All data are mean ± standard error.

### Aβ increases ODI, while effects of IL6 on ODI are strain-dependent

To further characterize spatial configuration of neurite structures, we performed analysis for ODI (Figure 6). Lower values of ODI are observed in large WM regions (which appear darker tones), such as splenium, corpus callosum, fimbria, and higher values are observed in grey matter areas (in various light grey tones). Consistent with an increase in NDI (presumed increased density of axonal/dendritic processes), we observed that Aβ increased their degree of orientation dispersion (geometric complexity or misorientation of axonal/dendritic processes). Thus, TgCRND8 had increased ODI in splenium, cerebral peduncle and hippocampus (splenium: main effect strain F_1,43_ = 33.8 p < 0.0001; cerebral peduncle: main effect strain F_1,43_ = 28.3 p < 0.0001; hippocampus: main effect strain F_1,43_ = 5.8 p = 0.01). The effects of IL6 were not widespread in the assessed structures and were also strain/tissue type dependent. Thus, in the cerebral peduncle (WM) IL6 reduced ODI only in Tg mice (treatment x strain interaction F_1,43_ = 12.4 p = 0.001 and main effect treatment F_1,43_ = 17.0 p = 0.0002), whereas in the hippocampus (grey matter) IL6 increased ODI only in nTg mice (main effect treatment F_1,43_ = 6.1 p = 0.02).

**Figure 6.**
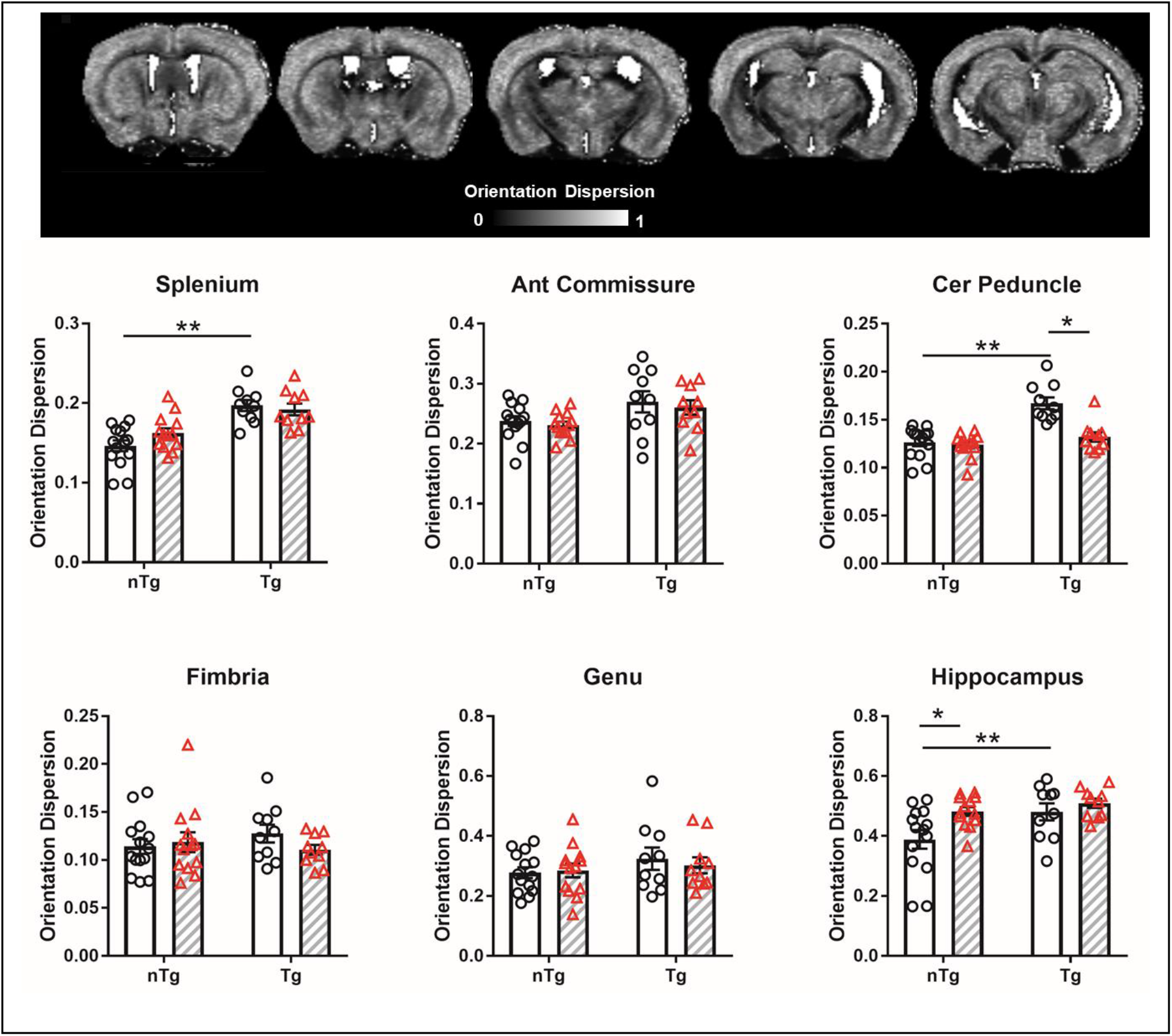
Aβ increases ODI in WM regions and hippocampus. The effects of IL6 differ in nTg and Tg mice. Representative ODI map shown. Clear bars (and circles) are control groups and hashed bars (and triangles) are IL6 treated. *significant difference between control and IL6 mice; **significant difference between Tg and nTg mice (Tukey’s post hoc test, p<0.05). All data are mean ± standard error.

### No effects of Aβ or IL6 on ISO index

The ISO index is part of the NODDI model that considers freely diffusing, unrestricted/unhindered water, that would be found mostly in cerebral spinal fluid (CSF) containing structures in the brain. As shown in Supplemental Figure 6, the greatest values of ISO are observed in ventricles. Darkest regions were found in grey matter and WM areas. Overall, we did not observe any effects of Aβ or IL6 on ISO values (Supplementary Figure 6).

### Aβ and IL6 alters a novel relationship between functional network connectivity and microstructure measures

We carried out functional MRI scanning at 11.1 Tesla to investigate if in addition to microstructural changes, IL6 or Aβ expression may affect intrinsic brain activity. For seed-based analysis, we chose cortical and hippocampal areas serving spatial and multi-sensory memory functions (Supplemental Figure 2). Statistical analysis using AFNI 3dMVM did not reveal any significant differences between the groups using this *a priori* seed-selection approach (data not shown). To supplement the seed-based functional connectivity approach, we conducted model free probabilistic ICA. ICA produced 30 components of which 19 were deemed qualitatively consistent with published resting state functional MRI networks in mice (Bukhari et al., 2017; Sforazzini et al., 2014; Shah et al., 2016). Figure 7A shows 9 components that included cerebellar, visual, retrosplenial, somatosensory, prefrontal, striatal, midbrain, hippocampal and thalamic regions. Figure 7B shows a comparison of component-24 for a network connecting thalamus, midbrain and anterior cingulate, which was similar in all 4 experimental groups. Permutation testing in FSL randomise revealed a significant main effect of Aβ on hippocampal-thalamic connectivity (component 24), and significant strain x treatment interaction on entorhinal-midbrain connectivity (component 21), striatum-parietal cortex connectivity (component 26) and striatum-cerebellum connectivity (component 26) (p < 0.05 familywise error corrected).

**Figure 7.**
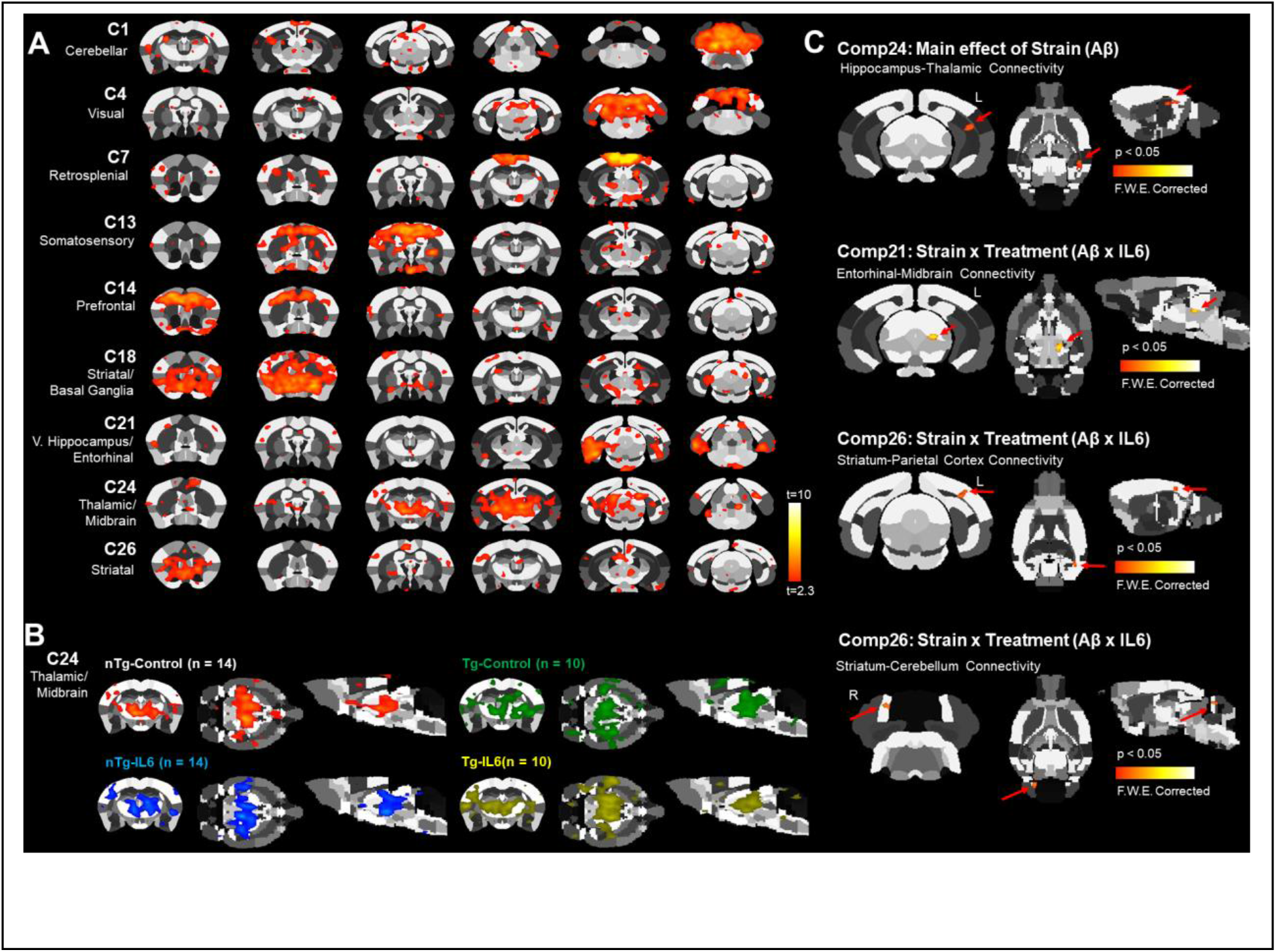
ICA of resting state fMRI reveals significant effects of Aβ and IL6 on hippocampal, thalamic, midbrain and striatal networks. A) Nineteen of 30 components were consistent with established ICA networks in mice. Shown here are 9, which include prefrontal, striatal, retrosplenial and hippocampal regions (t>2.3). B) Comparison of a thalamic/midbrain component in 4 experimental groups. C) Components that had significant main effects of Aβ or strain x treatment interactions following permutation tests (p<0.05 family-wise error corrected).

We additionally carried out graph theory analysis on a total of 4,005 pairwise comparisons between a total of 90 ROI’s (45 per mouse brain hemisphere) (Figure 8). Composite 3D functional connectome brains show similar node strengths and edge weights across the 4 experimental groups (network set at a density threshold to retain the top 2% of total edge weights) (Figure 8A). Composite matrices for 3D maps illustrate functional connectivity organizational patterns not observed after randomization of connections (null networks) (Figure 8A). TgCRND8 mice treated with IL6 had reduced modular organization than Tg controls (modularity: main effect treatment F_3,264_ = 3.7 p = 0.01). No additional differences were observed in clustering coefficient, small world coefficient, mean path length, or global node strength (Supplemental Figure 7).

**Figure 8.**
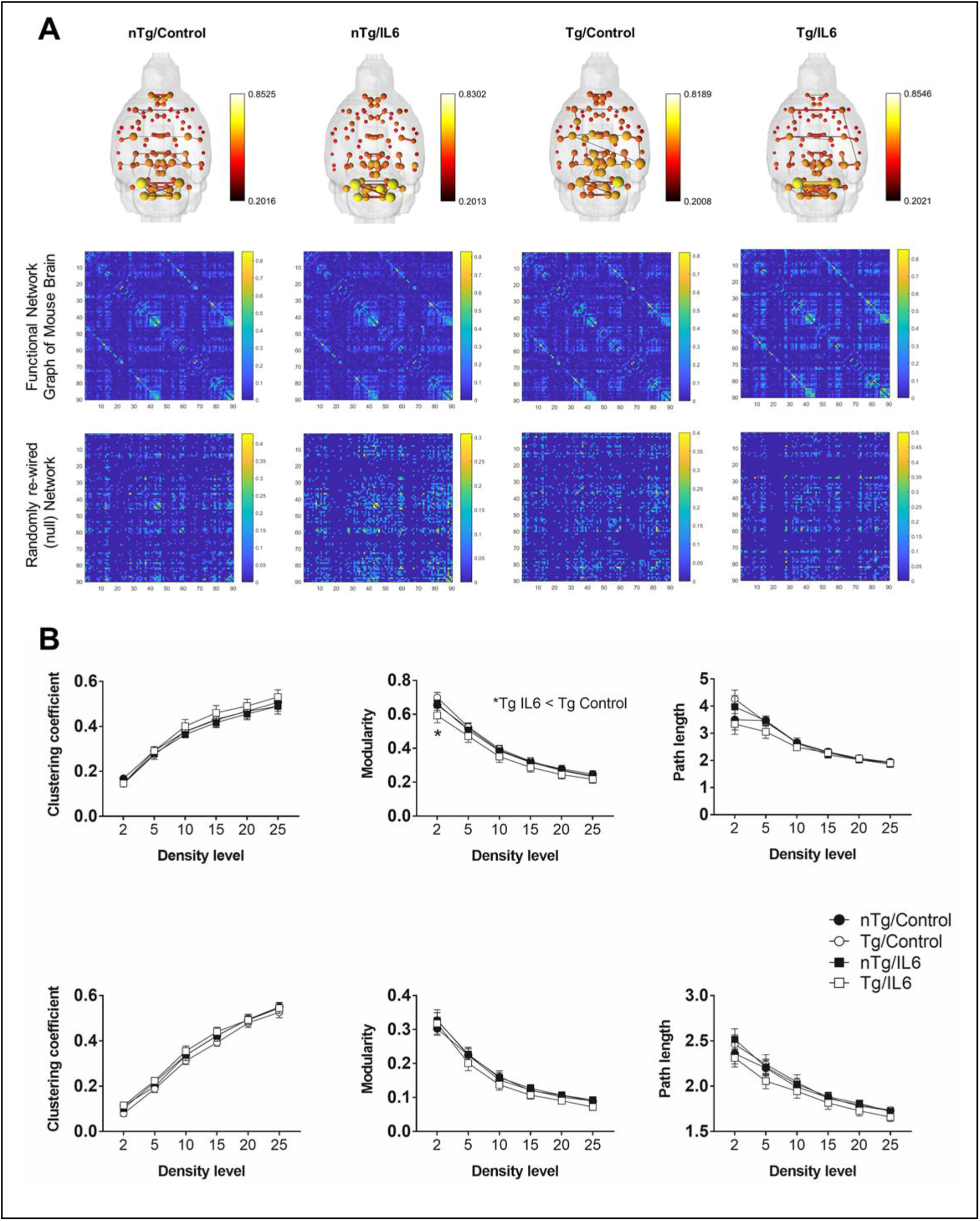
IL6 reduced modular organization but only in Tg mice. A) 3D functional connectome maps (density threshold set at 2%; map threshold at edge value of z=0.2). Below maps are composite graphs for each experimental group and their corresponding null networks. B) Connectomic metrics for graphs in A. Top row are functional connectome metrics and the bottom row show results for null networks. *significant difference between control and IL6 mice (Tukey’s post hoc test, p<0.05). All data are mean ± standard error.

Finally, to investigate the relationship between functional network metrics and DTI, free water and NODDI metrics we carried out correlation analyses. The first step involved normalization of dMRI metrics to their highest value and plotting these against modularity index, clustering coefficient, and node strength for each subject (Supplemental Figure 8). The results indicated that only clustering coefficient correlated significantly with dMRI metrics, specifically NODDI metrics. The results from non-normalized NODDI data suggest that clustering coefficient is altered by the degree of orientation dispersion and neurite density (Figure 9). Low orientation dispersion or neurite density correlates with high clustering. We observed this relationship in the splenium (NDI: p = 0.006; ODI: p = 0.04), hippocampus (NDI: p = 0.004; ODI: p = 0.004), and cerebral peduncle (NDI: p = 0.0008) (Figure 9) but found no significant correlations in genu and anterior commissure (data not shown). The fimbria showed a nonsignificant trend (NDI: p = 0.05). Inspection of individual data points revealed that IL6-treated nTg or TgCNRD8 mice (orange and green data points in plots in Figure 9) tended to have higher ODI and NDI values, which corresponded to lower clustering coefficient values than nTg control mice.

**Figure 9.**
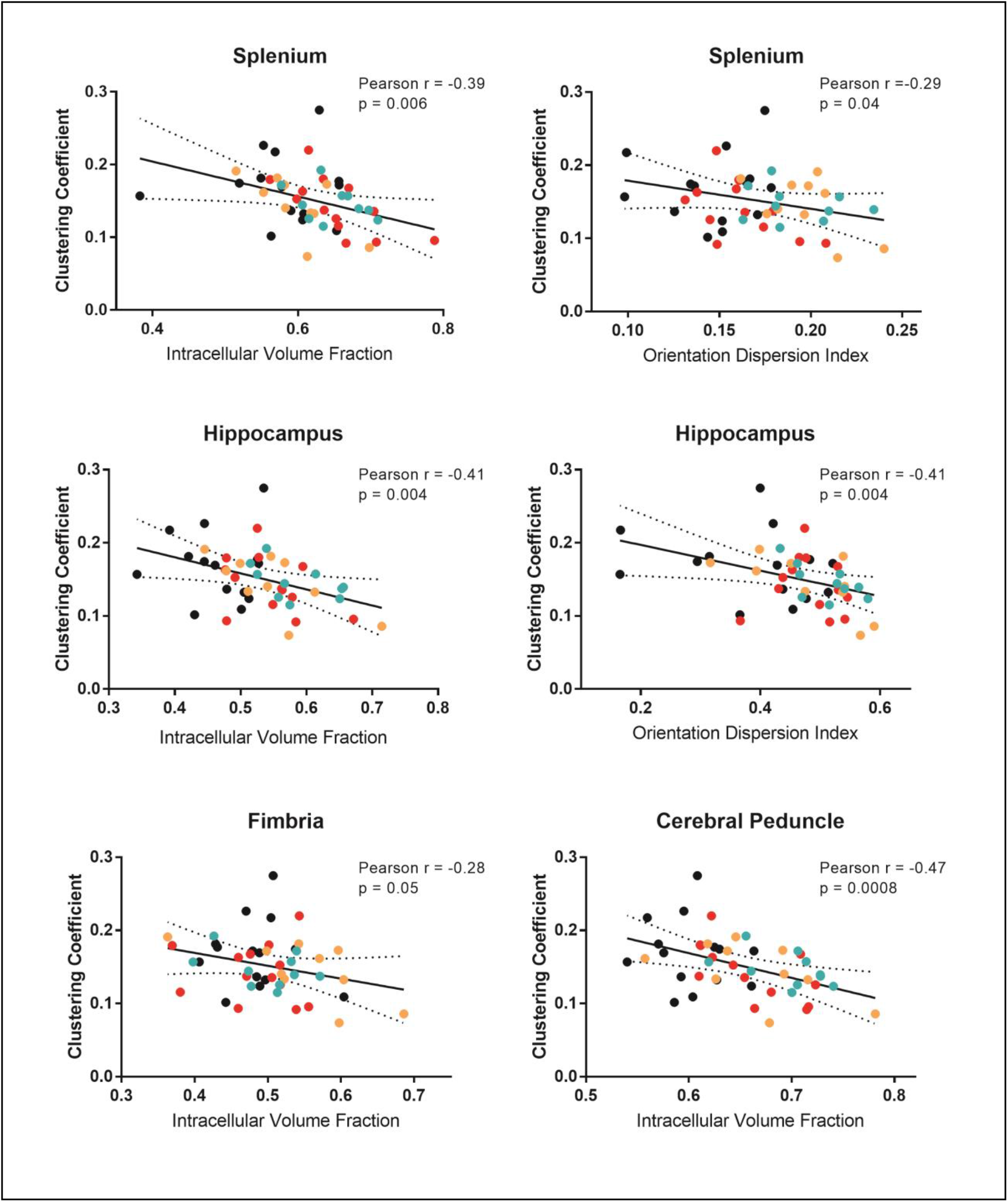
Aβ and IL6 influence the relationship between NODDI metrics and clustering coefficient. The degree of neurite density and orientation dispersion is negatively correlated with the clustering coefficient in splenium, hippocampus, and cerebral peduncle. Shown are individual data points for nTg-controls (black circles), nTg-IL6 (red circles), Tg-controls (orange circles), and Tg-IL6 (green circles). Statistical p values indicate a significant non-zero slope (99% confidence intervals shown).

## DISCUSSION

Using an Alzheimer’s disease mouse model, we carried out high field dMRI to investigate how amyloidosis with inflammatory conditioning induced by IL6 alters DTI, FW and NODDI metrics (summarized in Figure 10). Our data indicate that Aβ and IL6 differentially alter tissue microstructure measured by dMRI (e.g., fractional anisotropy, mean, axial and radial diffusivities, neurite density and orientation dispersion). Aβ and IL6 had diametrically distinct effects on FA and AD whereas their effects on free water and NODDI metrics were similar in directionality. Further, the effects of Aβ and IL6 on dMRI metrics were not additive, which argues against the possibility that Aβ and proinflammatory signaling through IL6 may constitute a double-hit in terms of exerting deleterious actions on tissue microstructure.

**Figure 10.**
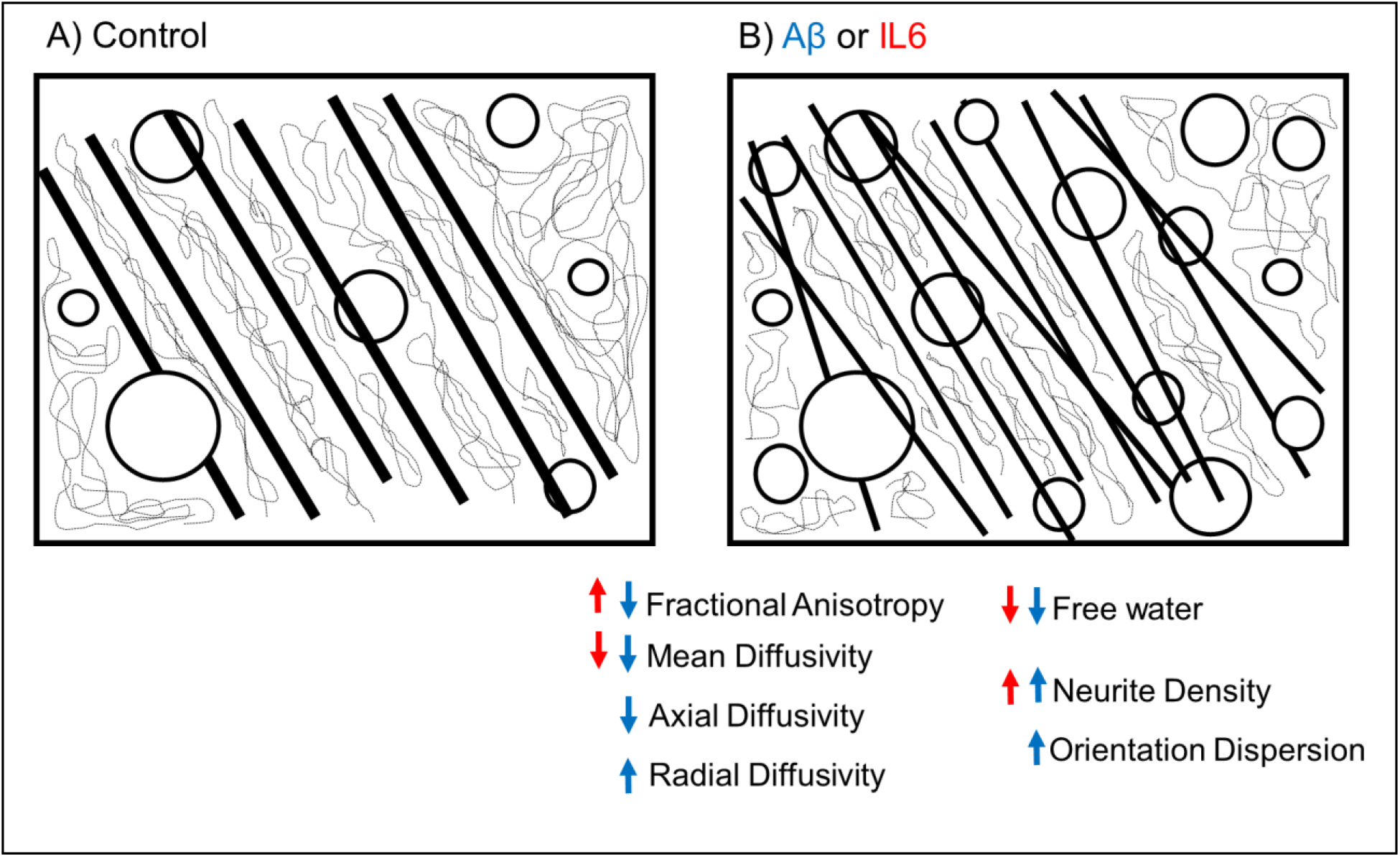
Summary schematic symbolizing microstructural changes contributing to the effects of Aβ or IL6 on DTI, free water and NODDI metrics. Schematic not meant to represent actual tissue microstructure but instead symbolize predicted alterations. A) Tissue microstructure conditions in a control state. B) Predicted microstructure conditions in the presence of Aβ or IL6. These may include axonal/dendritic atrophy with greater exchange across the membrane than along the length of processes (thinner lines in B vs A), increased presence of membranous structures (increased circles in B vs A), shorter diffusion paths (shorter hashed lines inside lines) and reduced extracellular volume for water movement (shorter hashed lines in the corners outside of lines), and finally, increased presence of misoriented axonal or dendritic processes (greater number of thinned lines at angles in B vs A). Blue arrows represent effects of Aβ and red the effects of IL6 on diffusion metric (increased or decreased effects).

We show that the TgCRND8 mice which shows early onset CNS amyloidosis had significant reductions in FA and MD values. This is consistent with previous dMRI research using TgCRND8 mice (Thiessen et al., 2010) and other APP mouse models (Mueggler et al., 2004; Sun et al., 2005), and our previous work using a frontal temporal dementia tauopathy model (Sahara et al., 2014). Reductions in these DTI metrics was associated with reduced AD (parallel diffusivity) and increased RD (perpendicular diffusivity) (Sun et al., 2005). The overall reduction of white matter AD might be associated with hindered diffusion along axons, which could be related increased concentrations of Aβ, other proteins involved in its production, or deficits in axonal transport (Johnson et al., 2010; Kim et al.; Zhang et al., 2009). Increased RD could be associated with increased membrane permeability, axon swelling, axon demyelination, or atrophy-like changes in dendritic processes (Adalbert et al., 2018; Zhang et al., 2017). Increased IL6 expression had mixed effects on tissue diffusivities and FA in nTg and TgCRND8 mice. The effect of IL6 was not synergistic with Aβ, and in the case of FA, IL6 appeared to counteract the effects of Aβ (Figure 2). This agrees with previous research suggesting protective actions of IL6 on Aβ burden in TgCRND8 mice (Chakrabarty et al., 2010b).

Interestingly, both Aβ and IL6 reduced free water levels (Figure 4). The free water method models unrestricted water mostly found in CSF, but also present in the extracellular space (Beaulieu, 2002), which can potentially be reduced in the presence of cells such as glia. Because IL6 and Aβ were independently observed to reduce free water relative to nTg controls, we predict that both proteins may be inducing an increase in glial cells in WM and hippocampal areas.

We also assessed NODDI metrics (Zhang et al., 2012) in the same animals processed for DTI and free water. The neurite density index (NDI) has been shown to correlate with histological markers of axonal or dendritic processes (Grussu et al., 2017), and changes in this NODDI metric has been associated with neurodegenerative changes (Colgan et al., 2016; Grussu et al., 2017; Parker et al., 2018). We observed that Aβ expression in TgCNRD8 mice was associated with increases in both NDI and the orientation dispersion index (ODI) in WM and hippocampus. At first glance this finding may seem counterintuitive considering the above-described reduced microstructural integrity. However, the results are consistent with previously reported negative correlations between FA and ODI (Zhang et al., 2012). Low FA values in WM correlate with high ODI, and this latter metric can show a positive correlation with NDI (Zhang et al., 2012). From a physiological standpoint, the co-occurrence of reduced microstructural integrity of axons along with increased density of misoriented axonal/dendritic processes in WM and hippocampus seems plausible given previously reported increases in activated microglial cells along with amyloid plaque-associated structural changes to neurons in mouse models with increased Aβ expression.

These findings in the TgCRND8 mouse are in partial agreement with previous NODDI results using the rTg4510 tauopathy mouse mode (Colgan et al., 2016). Colgan and colleagues reported increased MD and FA in hippocampus, reduced FA and increased MD in WM, and interestingly, reduced ODI and NDI in hippocampus and increased ODI and reduced NDI in WM. The rTg4510 mice were scanned at 8.5 mo when they display extensive intracellular tangle pathology in cortex and hippocampus, a loss of over 70% of pyramidal CA1 cells and dendritic spines (Santacruz et al., 2005). Further studies are needed to resolve differences in these two models of Alzheimer’s disease and related dementia hallmarks, particularly changes that occur with aging and their direct association with neural activity and cognitive behavioral impairments. As with the effects of IL6 on DTI and free water metrics, the effects of this inflammatory conditioning on NODDI measures varied. IL6 increased NDI in the splenium, anterior commissure, cerebral peduncle, genu, and hippocampus, but only in nTg mice. Similar to free water results summarized above, because Aβ and IL6 were independently observed to increase NDI and ODI relative to nTg controls, we predict that both these conditions (Aβ proteinopathy and IL6 inflammation) may be inducing similar microstructural changes to axons and dendritic processes in white matter and hippocampal areas.

As indicated in the introduction, Aβ and neuroinflammatory mechanisms overlap in the Alzheimer’s disease brain and their corresponding roles in pathological progression of dementia warrants investigation. Six-month old 5xFAD mice showed significant brain accumulation of activated microglia-specific protein tracer TSPO (Mirzaei et al., 2016). In PS2APP mice, there are significant increases in both Aβ and microglia levels from 5month to 16 months of age (Brendel et al., 2016). This is partially consistent with results in humans showing that in advanced Alzheimer’s disease there is a close relationship between amyloid burden with neuroinflammation (Kreisl et al., 2013). However, non-Alzheimer’s disease individuals with mild cognitive impairment (MCI) either fail to show significant microgliosis concomitant with regions of high amyloid burden (Kreisl et al., 2013), or only a subset of MCI subjects show overlapping increases in amyloid and neuroinflammation (Parbo et al., 2017). A longitudinal study identified a transient elevation in microglia early in MCI which then peaks again later in Alzheimer’s disease (Fan et al., 2017). Moreover, areas showing high plaque load do not necessarily show increased activated microglia (Serriere et al., 2015). It is possible that the functional and structural outcome of CNS immune responses depend on the specific cytokine-initiated immune response, with some inflammatory mechanisms exacerbating neurodegenerative phenotypes and Aβ deposition while others serving protective roles (Chakrabarty et al., 2010a; Chakrabarty et al., 2011a; Chakrabarty et al., 2011b; Chakrabarty et al., 2010b; Chakrabarty et al., 2012). Comparisons of DTI, free water, and NODDI measures after treatment with these different cytokines will be important to understanding the link between dMRI metrics and CNS immunity.

Our results are for the most part in agreement with prior studies using APP mouse models. No changes in the apparent diffusion coefficient (ADC) were observed when comparing young 3 mo to old 12-16 mo TgCNRD8 mice (Thiessen et al., 2010), although reduced ADC were observed in grey matter areas of 25 mo compared to 6 mo APP23 mice (Mueggler et al., 2004). Reductions and increases in FA were measured in 23 mo Tg2576 APP mice scanned at 11.1 Tesla, and this involved reduced MD, AD, and RD (Muller et al., 2013). In APP/PS1 mice, Qin et al measured increased FA, AD, and RD in cortex, dorsal striatum, thalamus, and in WM structures (Qin et al.), and their electron microscopy results indicated widespread swelling of axons, hypertrophic astrocytes and neuronal loss (Qin et al.). The latter results were highly consistent with those of Shu et al (2013)(Shu et al., 2013). In the triple transgenic (3xTgFAD) mouse model, reduced FA and AD was measured only in the hippocampus (Snow et al., 2017). Results similar to those obtained here were reported by Sun et al, who observed reduced relative anisotropy along with reduced AD and increased RD in the Tg2576 mouse model (Sun et al., 2005). Interestingly, opposite changes in hippocampal MD, AD and RD was observed between 6-8 mo in the APP/PS1 mouse using a diffusion kurtosis image acquisition scheme (Praet et al., 2018). The varied results across studies illustrates difficulties inherent to *in vivo* dMRI of Alzheimer’s disease mouse models, which setting aside technical challenges, may be influenced by the dynamic nature of the cellular and microstructural changes occurring in response to the progression of AD.

Finally, we observed an inverse linear relationship between NDI, ODI in WM and hippocampus and the functional connectomic clustering coefficient. High NDI and ODI was negatively correlated with low clustering, which suggests that increasing the density and orientation dispersion of neurites may diminish functional network organization in the brain. Implied in this relationship is that in addition to capturing neuronal fiber organization, NODDI metrics somehow measures components of microstructure that may describe or relate to neuronal activity and how neuronal communication is organized in the brain. Recently it was reported that high neurite density in auditory/temporal cortex in humans was associated with a decrease latency (faster processing) of neurophysiological processing of speech (Ocklenburg et al., 2018). Also recently, Brown and colleagues observed that decreased default mode network (DMN) deactivation during a working memory task was associated with Alzheimer’s disease pathology and cognitive decline (Brown et al., 2018). Interestingly, decline in WM integrity was reported to influence the reduced DMN deactivation, suggesting a relationship between microstructural differences between controls and Alzheimer’s disease subjects and network activity. Our results suggest a negative relationship potentially involving reduced activity or reduced modular organization in Alzheimer’s disease mice and with IL6 treatment, which is contrary to the positive relationship reported by Ocklenberg et al (Ocklenburg et al., 2018) and Brown et al (Brown et al., 2018). A possibility is that the increased NDI and ODI in the present work, and its association with reduced clustering, might involve increased presence of neuroglial cells (as noted above) and not necessarily changes in the density of functionally active axons and dendritic processes. Moreover, the fact that both NDI and ODI were increased may suggest atrophic microstructural changes in TgCRND8 and IL6 treated mice that may adversely impact neural processing. It will be interesting to resolve such interactions between these two different imaging modalities. The relationship between NODDI and graph theory metrics warrants further investigation in the development of novel biomarkers for Alzheimer’s disease and other neurodegenerative diseases.

## Acknowledgements

The authors thank the Advanced Magnetic Resonance Imaging and Spectroscopy (AMRIS) facility and National High Magnetic Field Laboratory (NHMFL) for their continued support (National Science Foundation Cooperative Agreement No. DMR-1157490 and the State of Florida).

