## Supplemental Data for "Increased Neurite Orientation-Dispersion and Density in the TgCRND8 Mouse Model of Amyloidosis: Inverse Relation with Functional Connectome Clustering and Modulation by Interleukin-6"

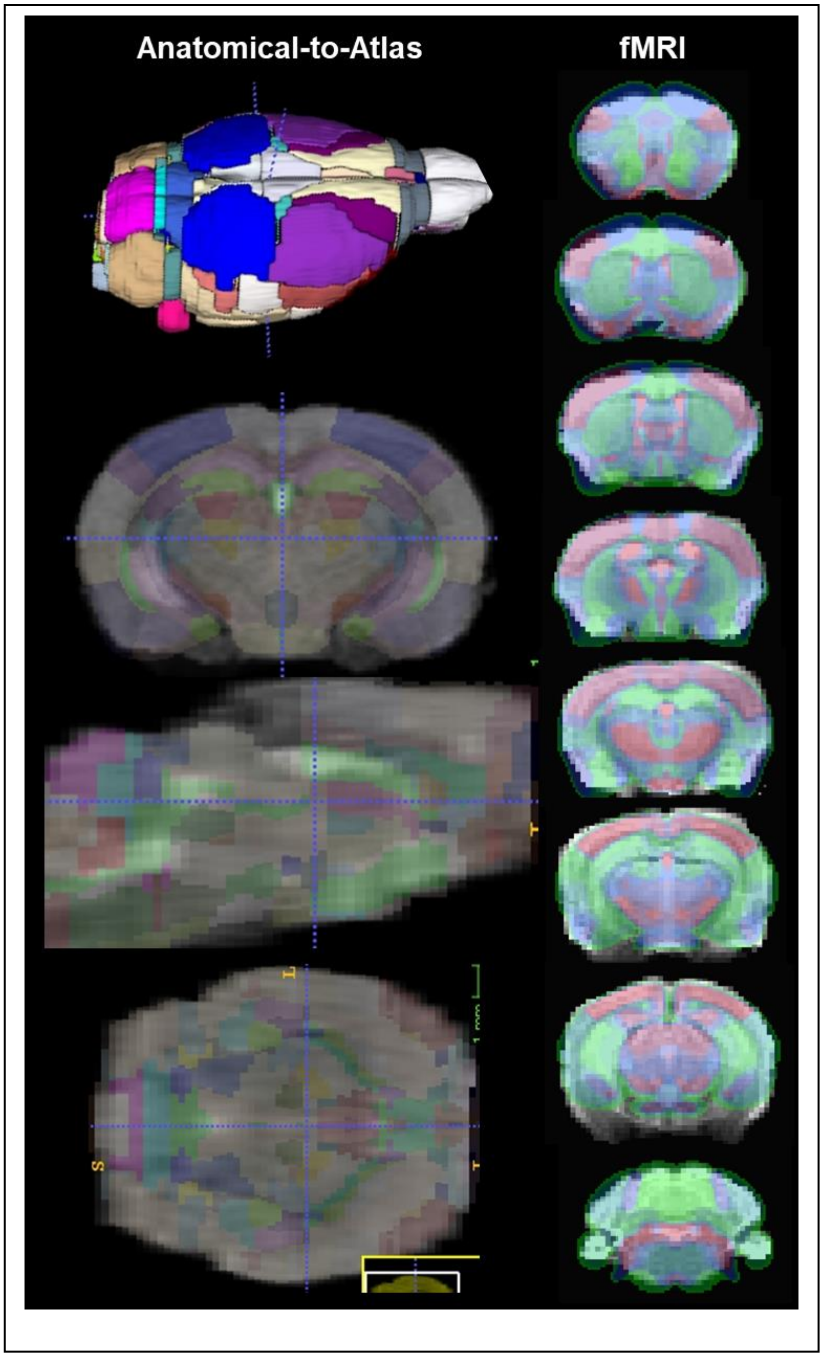


**Supplemental Figure 1.** Representative atlas-aligned T2 Turbo RARE image and functional MRI scan.

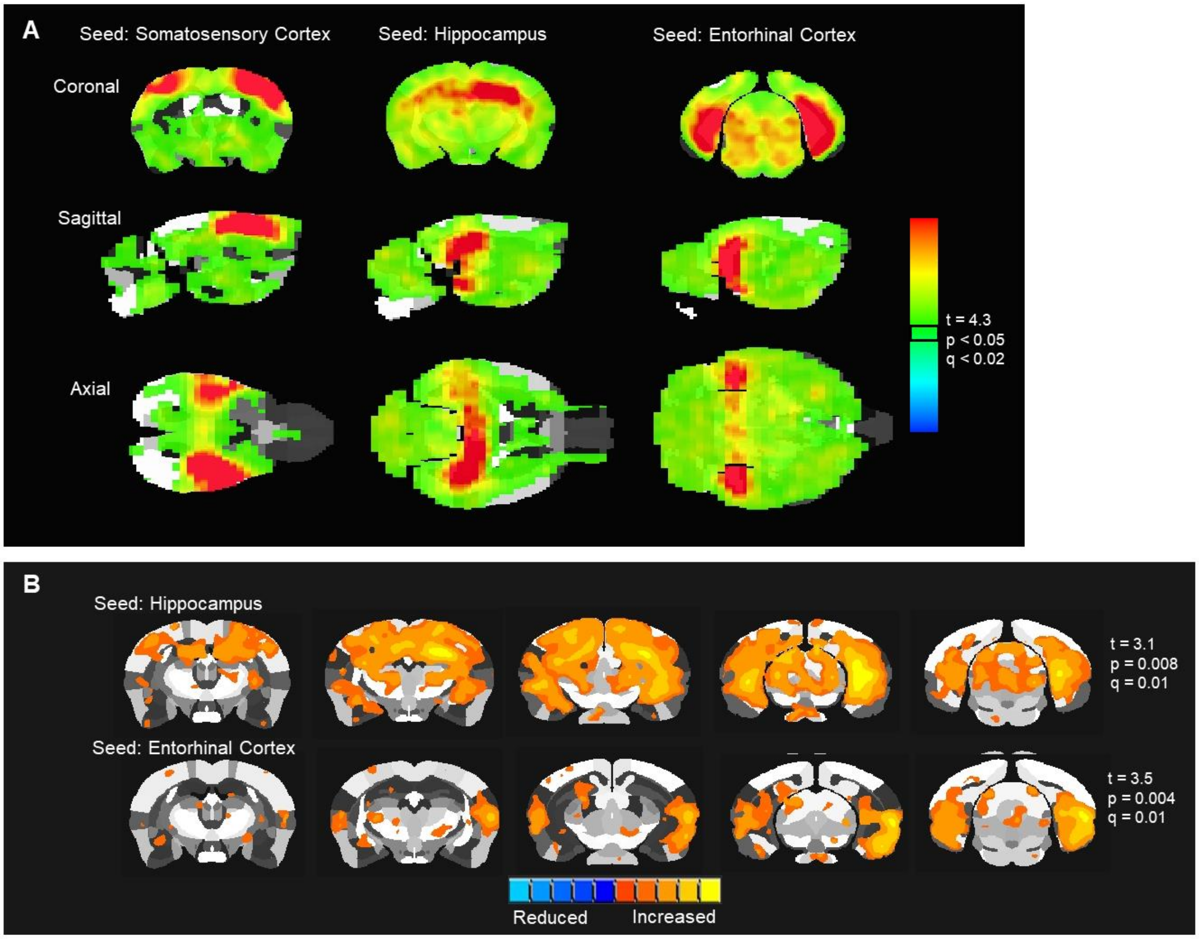


**Supplemental Figure 2.** Composite statistical maps of seed-based functional connectivity for hippocampus, entorhinal and somatosensory cortex. A) Statistical maps of 48 mouse brains with seed placements indicated above maps. Maps are shown in 3 different views. Scale bar indicates significance level (threshold at t=4.3, FDRq<0.02 and p<0.05). B) Statistical maps for nTg control mice (n=14) for hippocampus and entorhinal seeds.

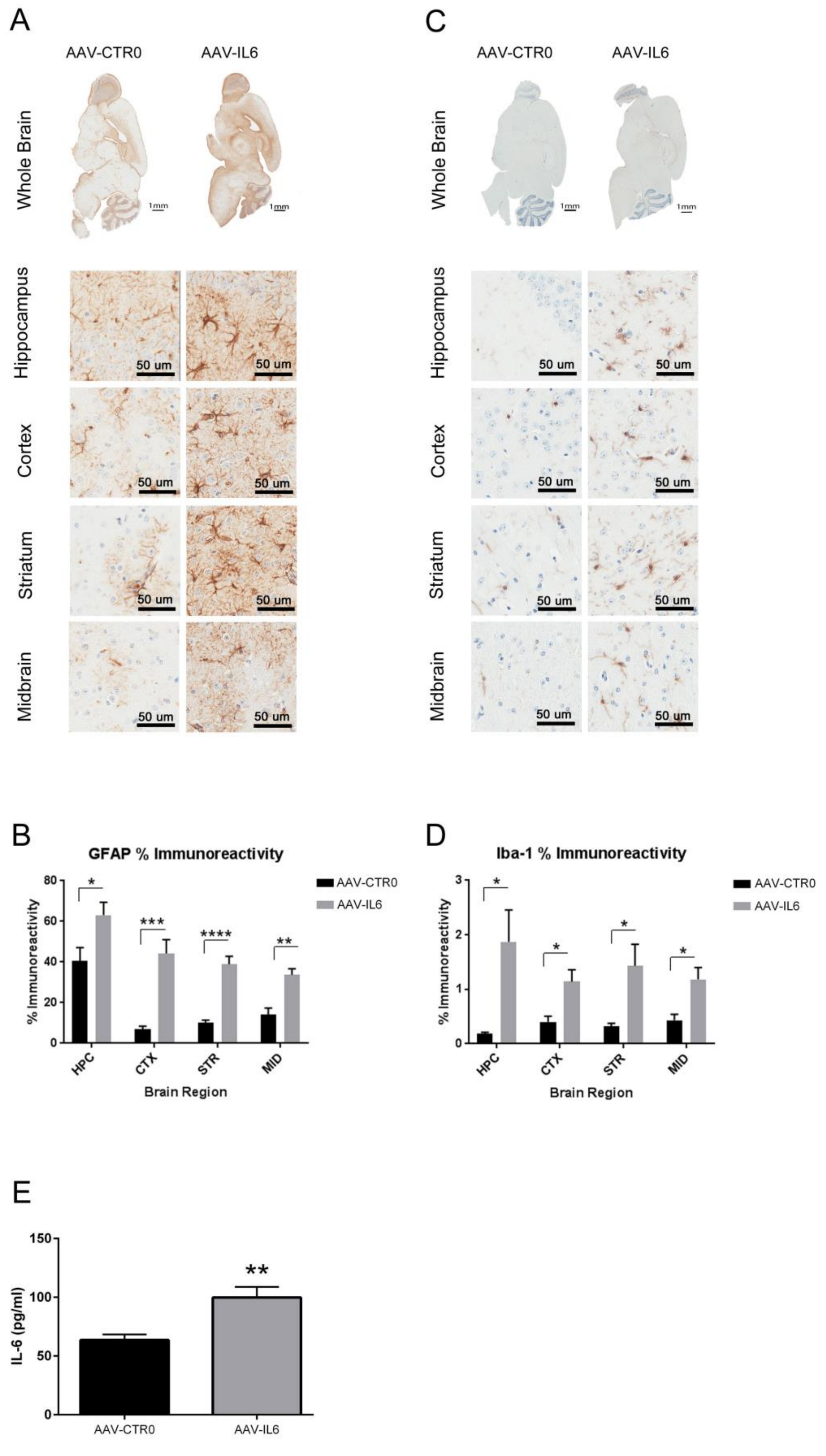
A
AAV-CTR0
AAV-IL6
Whole Brain
1mm
Hippocampus
Cortex
Striatum
Midbrain
50 um
C
AAV-CTR0
AAV-IL6
B
GFAP % Immunoreactivity
% Immunoreactivity
80
60
40
20
0
HPC
CTX
STR
MID
Brain Region
*
***
****
**
AAV-CTR0
AAV-IL6
D
Iba-1 % Immunoreactivity
% Immunoreactivity
3
2
1
0
HPC
CTX
STR
MID
Brain Region
E
IL-6 (pg/ml)
150
100
50
0
**
AAV-CTR0
AAV-IL6

**Supplemental Figure 3.** CNS expression of IL6 results in widespread gliosis. A) Analysis of astrocytosis (GFAP, A-B), microgliosis (Iba-1, C-D) and IL6 protein (E) in nTg mice demonstrate that IL6 expression leads to dramatic upregulation of immune response throughout the brain. HPC, hippocampus; CTX, cortex; STR, Striatum; MID; midbrain. B-D. One way ANOVA, *p<0.05; **p<0.01, ***p<0.005, n=6; E. Student's t test, **p<0.01, n=4.

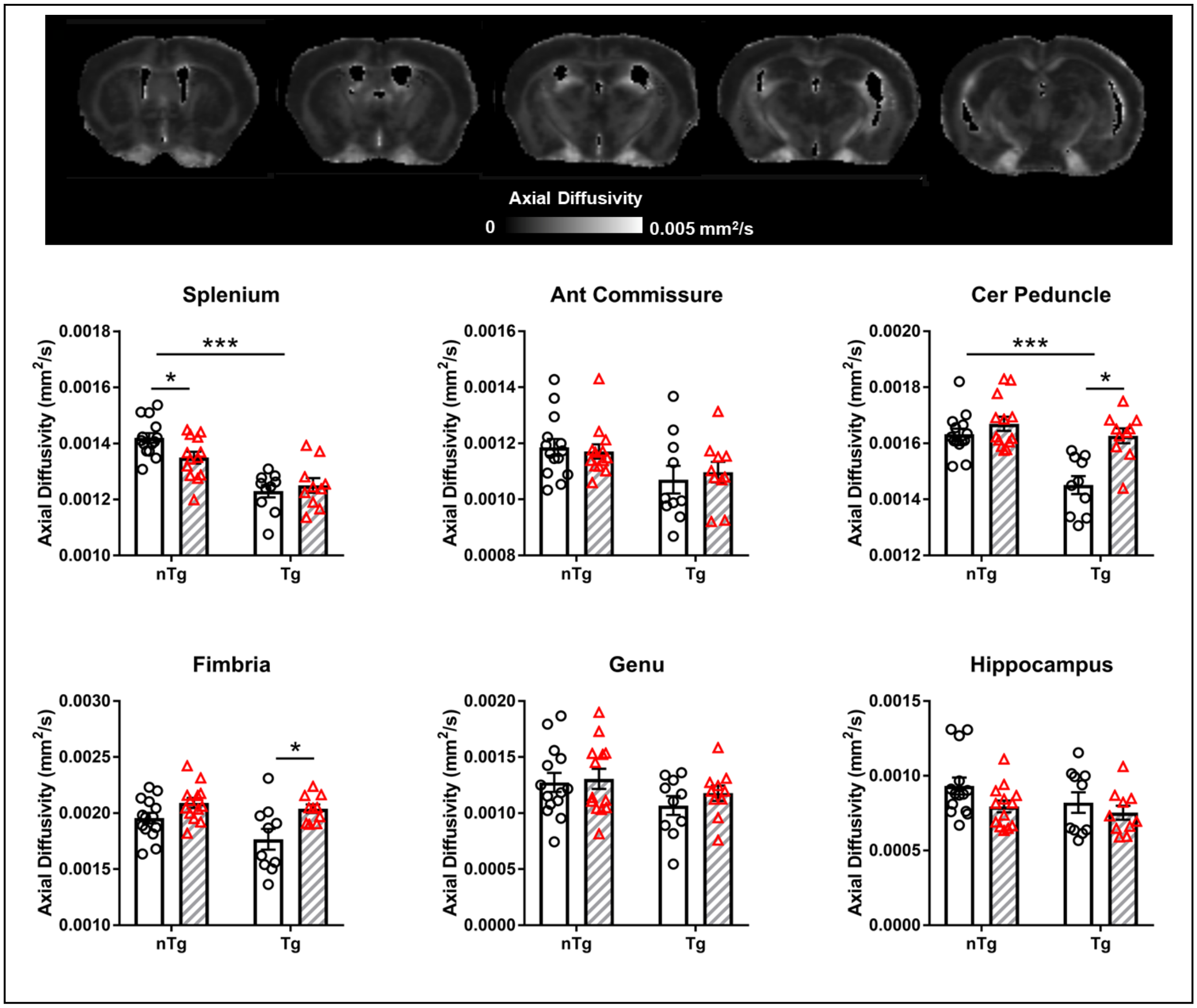


**Supplemental Figure 4.** Aβ reduces AD in WM regions and the effects of IL6 on AD are strain dependent. Representative AD map shown. Clear bars (and circles) are control groups and hashed bars (and triangles) are IL6 treated. *significant difference between control and IL6 mice; ***significant difference between all Tg and all nTg mice (Tukey's post hoc test, $p<0.05$).

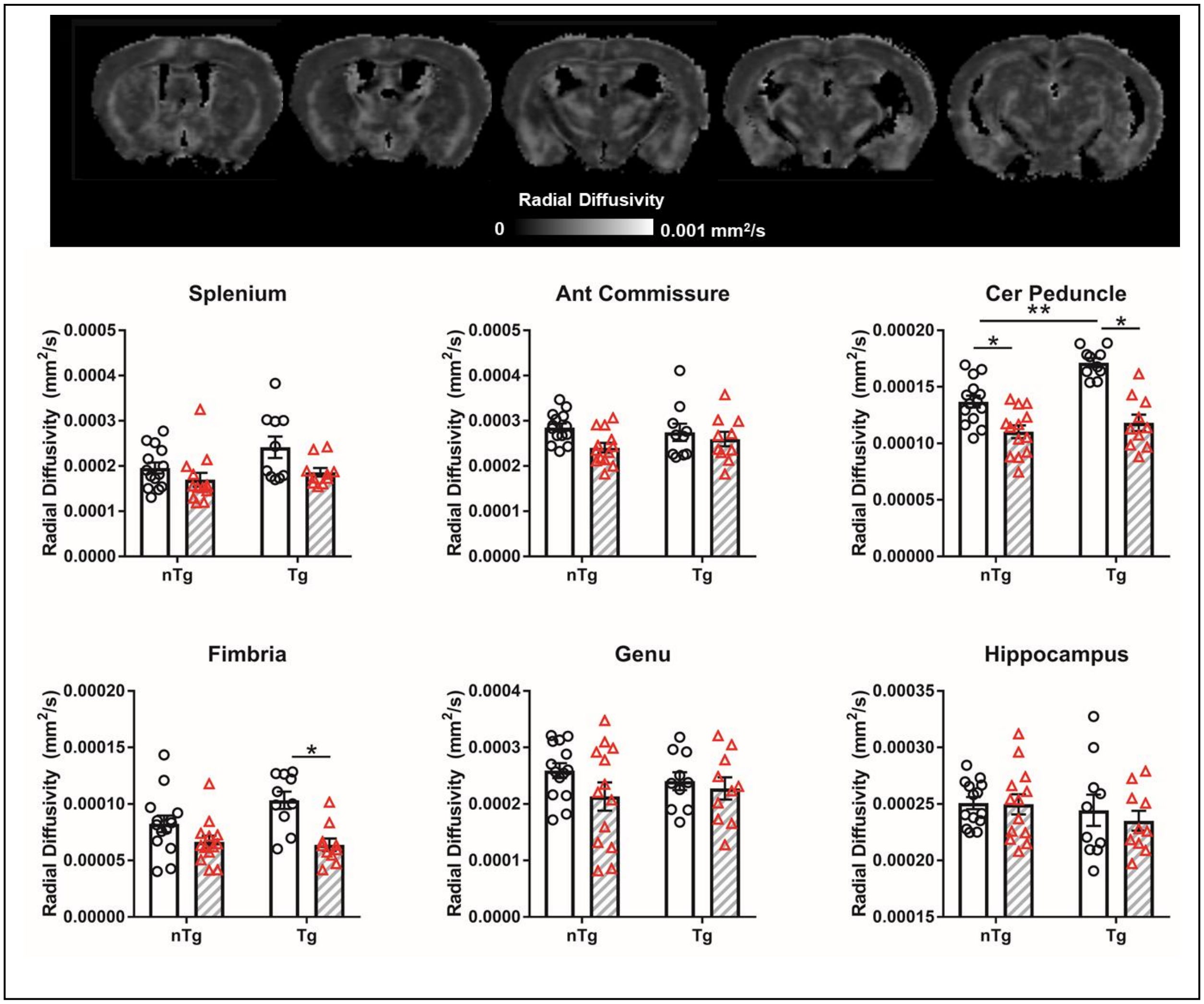


**Supplemental Figure 5.** Aβ increases RD in cerebral peduncle and IL6 reduces RD. Representative RD map shown. Clear bars (and circles) are control groups and hashed bars (and triangles) are IL6 treated. *significant difference between control and IL6 mice; **significant difference between Tg and nTg mice (Tukey's post hoc test, $p<0.05$).

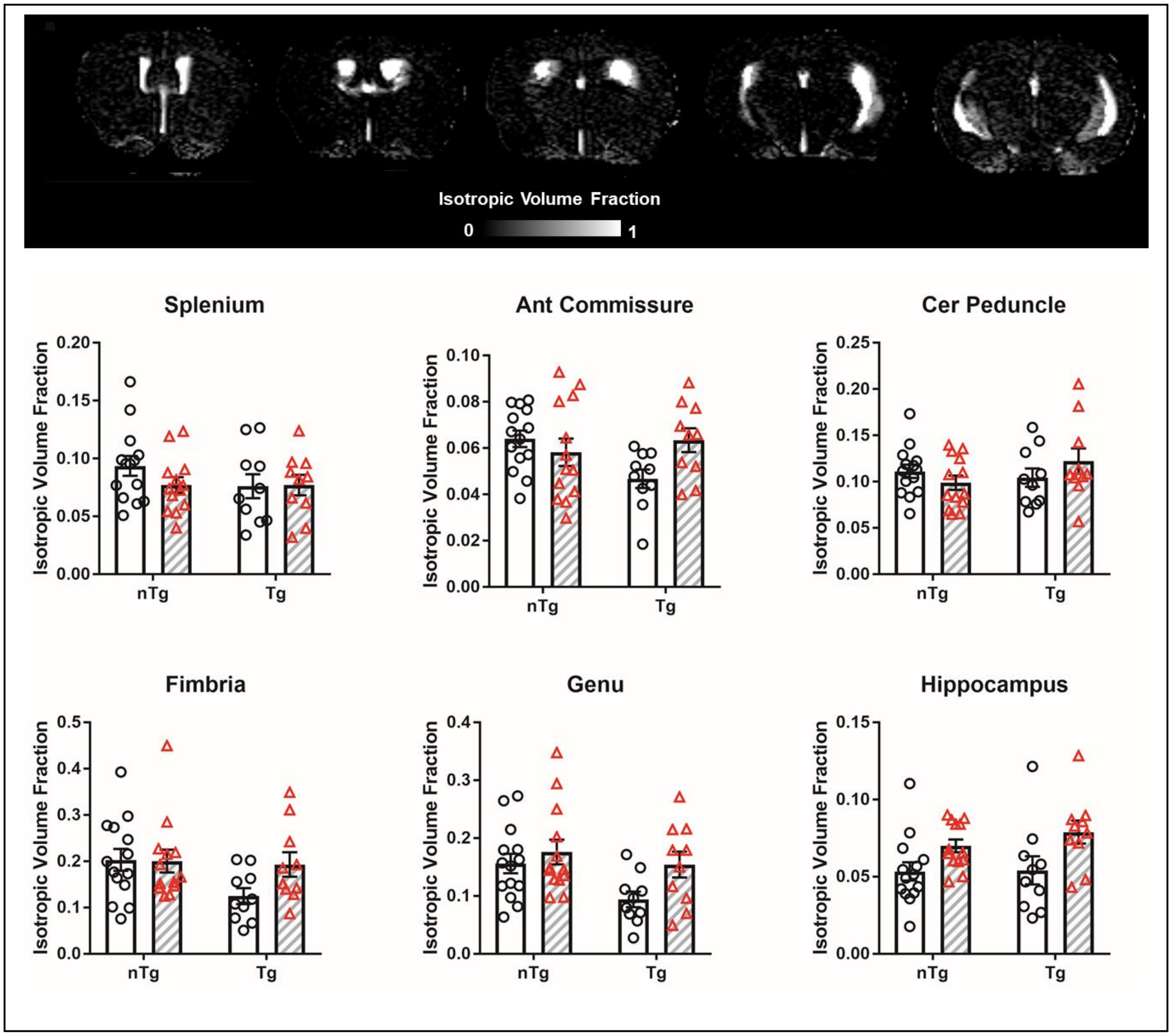


**Supplemental Figure 6.** No effect of Aβ or IL6 on ISO index. Representative ISO map shown. Clear bars (and circles) are control groups and hashed bars (and triangles) are IL6 treated. *significant difference between control and IL6 mice.

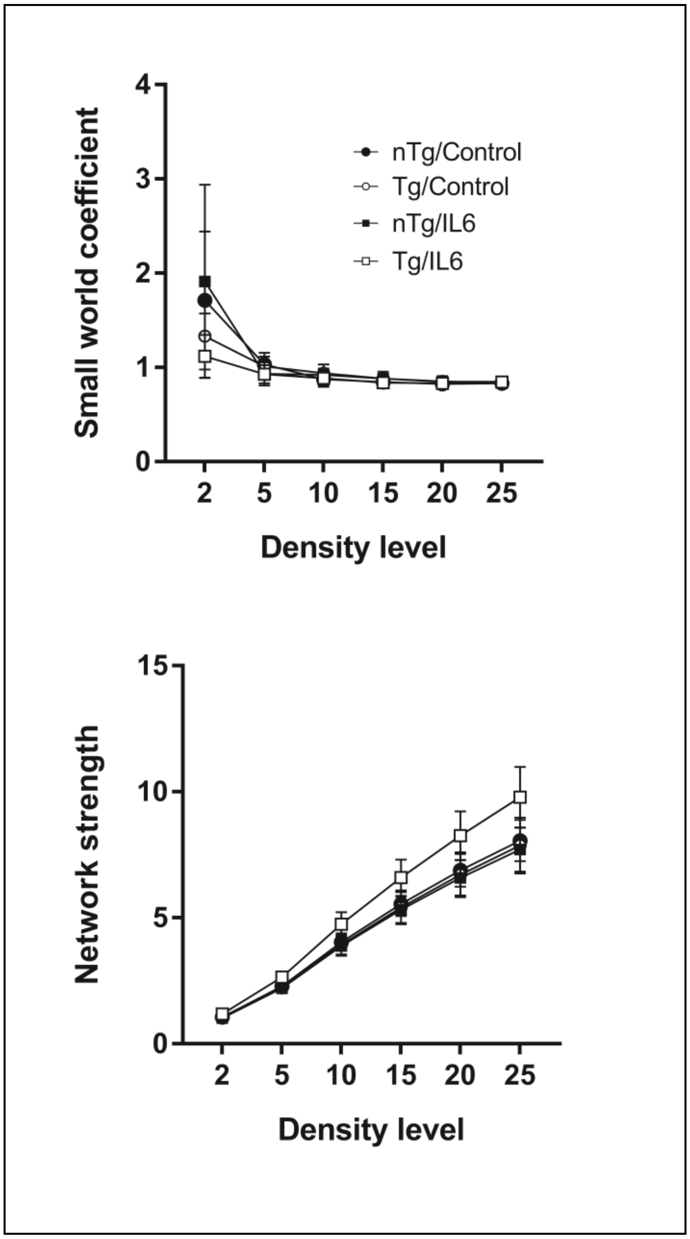


**Supplemental Figure 7.** No effect of Aβ or IL6 on small world coefficient and node strength. All data are mean ± standard error.

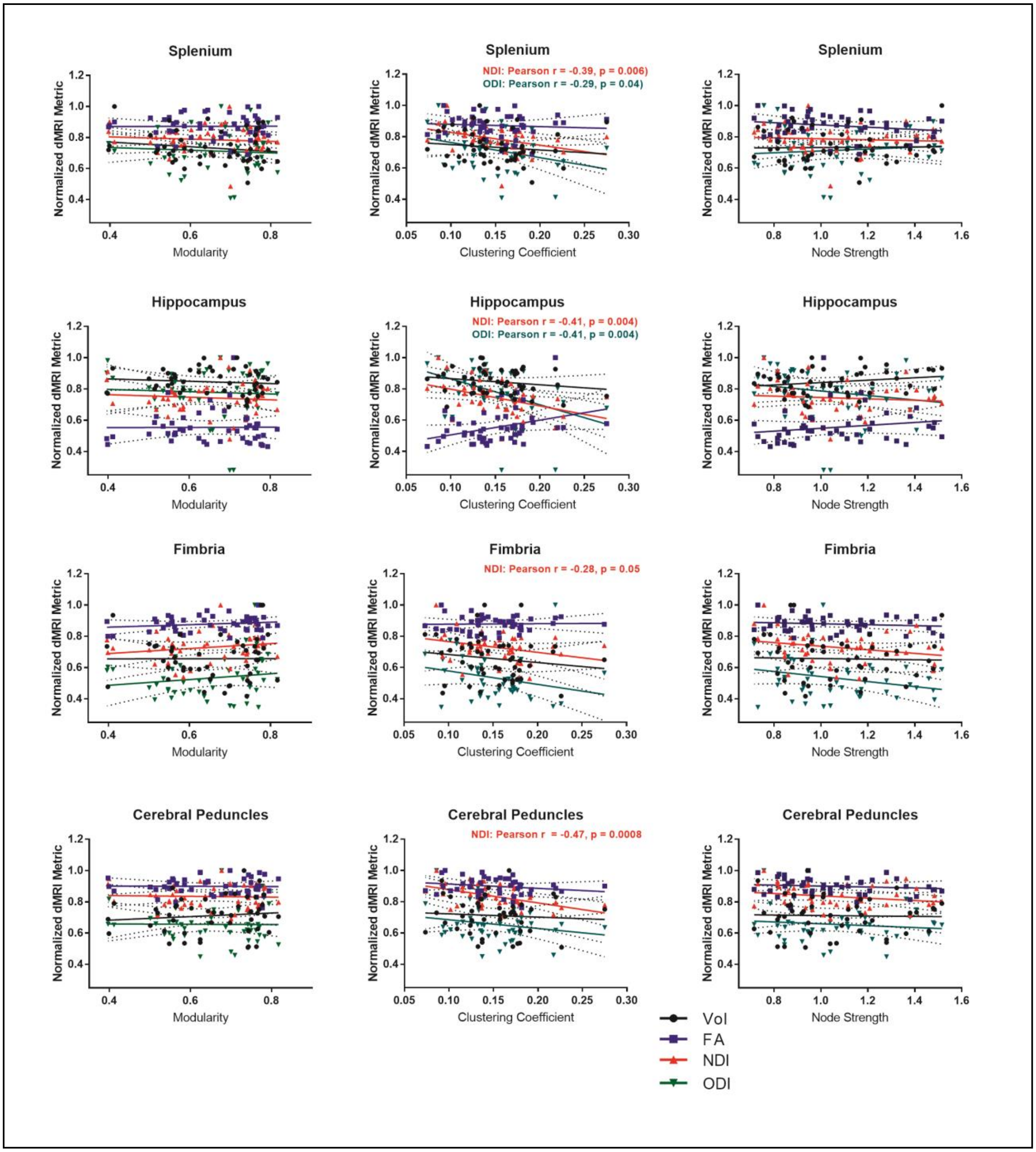


**Supplemental Figure 8.** Aβ and IL6 influence the relationship between NODDI metrics and clustering coefficient. This is observed in splenium, hippocampus, and cerebral peduncle, but not genu or anterior commissure (fimbria shows non-significant trend), nor with ROI volume. Modularity and node strength did not show significant correlations with diffusion MRI metrics or ROI volume. Coloring of data points and regression lines indicates ROI volume (black), fractional

FA (blue), NDI (red), and ODI (green). Statistical p values indicate a significant non-zero slope (99% confidence intervals shown).
